# Three-dimensional Genome Structure Reveals Distinct Chromatin Signatures of Mouse Female Germline Stem Cells During Development

**DOI:** 10.1101/787689

**Authors:** Geng G. Tian, Xinyan Zhao, Wenhai Xie, Xiaoyong Li, Changliang Hou, Yinjuan Wang, Lijuan Wang, Xiaodong Zhao, Hua Li, Jing Li, Ji Wu

## Abstract

The three-dimensional configuration of the genome ensures cell-type-specific gene expression profiles by placing genes and regulatory elements in close spatial proximity. Here, we revealed the distinct features of the chromatin architecture in female germline stem cells (FGSCs) by *in situ* high-throughput chromosome conformation analysis. We also showed that the X chromosome structures were similar in spermatogonial stem cells and FGSCs. Using integrative analysis of the three-dimensional chromatin structure, we observed that the TADs were attenuated in germinal vesicle oocytes and disappeared in metaphase II oocytes during FGSC development. Finally, we identified conserved compartments belonging to the paternal/maternal genomes during early embryonic development, which were related to imprinted genes. These results will provide a valuable resource for studying and further our understanding of the fundamental characteristics of oogenesis and early embryo development.

## INTRODUCTION

The chromatin architecture of germline stem cells (GSCs) carries the information necessary for the cells to exert their unique functions, and is thus an essential factor in the transmission of the genome from generation to generation. GSCs can renew themselves and differentiate into gametes, including sperm and metaphase II (MII) oocytes^1–3^. During this process, spermatogonial stem cells (SSCs) differentiate into sperm by packaging the chromatin into a highly condensed configuration. Recent identified female GSCs (FGSCs) in postnatal ovaries were shown to differentiate into MII oocytes after transplantation into the ovaries of infertile mice^4–10^, thus reshaping the idea that female mammals lose the ability to produce oocytes at birth^11,12^ Unlike other stem cells, GSCs can undergo meiosis to produce haploid gametes with chromatin remodeling. It is therefore necessary to characterize the chromatin structure of GSCs during their development to further our understanding of GSC biology.

High-throughput chromosome conformation (Hi-C) is a powerful technology for studying the genome-wide architecture, allowing the high-order chromatin structure to be displayed and revealing the chromatin organization in the nucleus^13^. The spatial organization of chromatin, as the structural and functional basis of the genome, can affect DNA localization, with important roles in gene transcription, the prevention of DNA damage, and ensuring DNA duplication and other biological processes^14,15^. Previous studies reported that the chromatin architecture changed dynamically during spermatogenesis, with dissolved and reappeared topologically associated domains (TADs) and A/B compartments^16,17^ However, the signature of the chromatin architecture during the development of FGSCs is unknown. Furthermore, the paternal and maternal chromatin architectures have been shown differences during early embryonic development^18,19^, but the respective changes in high-order allelic genome structure in early embryonic development remain to be explored.

In this study, we used *in situ* Hi-C technology to compared the chromatin organization of FGSCs with pluripotent stem cells (induced pluripotent stem cells, iPSCs), adult stem cells (ASCs) including SSCs and neural stem cells (NSCs), and somatic cells (mouse SIM embryonic fibroblast cells, STOs) to explore the chromosome structure character of FGSCs. Together with RNA sequencing (RNA-Seq) and chromatin immunoprecipitation sequencing (ChIP-Seq), we identified the distinct features of chromatin organization in FGSCs at three major levels: A/B compartments, TADs, and chromatin loops, and demonstrated that FGSCs were most similar to other ASCs, and largely different from iPSCs and STOs. We also identified similarities in X chromosomes between SSCs and FGSCs by principal component analysis of the X chromosome. Further analysis of Hi-C data during female germline cell development showed that TADs were attenuated but still present in germinal vesicle (GV) oocytes, disappeared in MII oocytes. Finally, we found conserved compartment regions in the paternal/maternal genomes in early embryonic development, related to imprinted genes. Together, these findings revealed the unique chromatin signature of FGSCs, and presented a whole landscape of high-order genome structure during development of female germline cells and early embryos.

## RESULTS

### Biological Characterization of FGSCs and Other ASCs

FGSCs were isolated and cultured from the ovaries of Ddx4-Cre;mT/mG neonatal mice, as described previously^20^. After culture for at least 18 passages, the cells exhibited a characteristic morphology similar to that previously described for FGSCs^2,20^. The expression of female germline marker genes was determined by reverse-transcription polymerase chain reaction (RT-PCR). FGSCs after long-term culture expressed *Oct4, Fragilis, Mvh* (mouse vasa homologue, expressed exclusively in germ cells), *Stella, Gfrα1*, and *Dazl* genes. Furthermore, immunofluorescence analysis revealed that these cells also expressed MVH, confirming their identity as FGSCs (Figure 1A).

**Figure 1.**
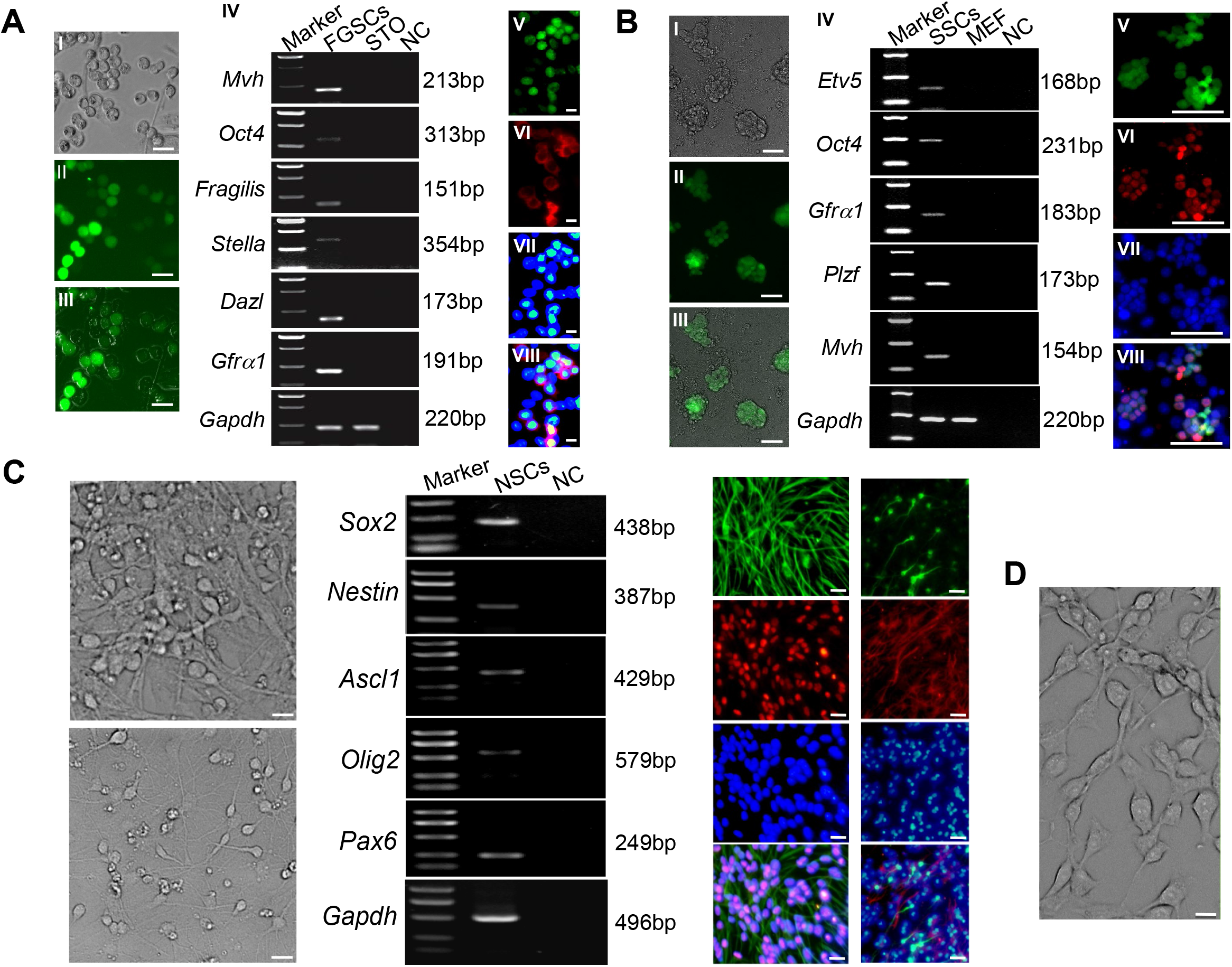
Morphology and biological characteristics of FGSCs and other ASCs. (A) Morphology and biological characteristics of FGSCs. (I) Representative morphology of cultured FGSCs (>18 passages) from Ddx4-Cre;mT/mG mice. (II) Representative field under UV light for FGSCs with GFP expression. (III) Merged images from I and II. Bar=25 μm. (IV) RT-PCR analysis of female germline markers. Marker, 100-bp DNA markers. (V–VIII) Immunofluorescence of FGSCs for GFP (V), MVH (VI), DAPI (VII), and merged (VIII). Bar=10 μm. (B) Morphology and biological characteristics of SSCs. (I) Representative view of cultured SSCs (>20 passages) from CAG-EGFP mice. (II) Representative image under UV light for SSCs with GFP expression. (III) Merged images from I and II. Bar=25 μm. (IV) RT-PCR analysis of male germline markers. Marker, 100-bp DNA markers. (V–VIII) Immunofluorescence of SSCs for GFP (V), PLZF (VI), DAPI (VII), and merged (VIII). Bar=50 μm. (C) Morphology and biological characteristics of NSCs. (I) Representative morphology of cultured NSCs. (II) Representative view of differentiated NSCs *in vitro*. Bar=10 μm. (III) RT-PCR analysis of NSC markers. Marker, 100-bp DNA markers. (V–VIII) Immunofluorescence of NSCs for Nestin (V), SOX2 (VI), DAPI (VII), and merged (VIII). (IX–XII) Immunofluorescence of differentiated NCs for TUJ-1 (IX), GFAP (X), DAPI (XI), and merged (XII). Bar=10 μm. (D) Representative morphology of cultured STOs. Bar=10 μm.

We isolated and cultured SSCs from the testes of 6-day-old DBA/2×CAG-EGFP F1 mice, as described previously^21^. Long-term cultured SSCs (>20 passages) were assessed by RT-PCR and were shown to express male germline marker genes (*Etv5, Oct4, Plzf* (promyelocytic leukaemia zinc finger), *Gfrα1*, and *Mvh*). These results were confirmed by immunofluorescence and most of the cultured cells were also positive for PLZF expression, confirming their identity as SSCs (Figure 1B).

Isolated primary NSCs could self-proliferate and be cultured for 5–8 passages in NSC proliferation medium. Morphologically, cultured NSCs were spindle-shaped with a high nucleus-to-cytoplasm ratio, as reported previously^22^ The cultured NSCs were positive for several NSC-specific markers including *Nestin, Sox2, Pax6, Olig2*, and *Ascl1*, as determined by RT-PCR. Immunocytochemical staining confirmed that most of the cultured NSCs were positive for NESTIN and SOX2, as typical markers specific for NSCs. Following the removal of epidermal growth factor and basic fibroblast growth factor from the medium, NSCs spontaneously differentiated into neurons and astrocytes, characterized by prominent dendrites with long axons and by extensive cytoplasm with thick processes, respectively. The differentiation potential of cultured NSCs was confirmed by immunochemical staining of the neural- and astrocyte-specific markers TUJ-1 (β3 Tubulin) and GFAP (glial fibrillary acidic protein). These results confirmed the identity of the cultured cells as NSCs (Figure 1C). The morphology of STOs is shown in Figure 1D.

### Global Chromosome Organization Map in FGSCs

To study the signature of the chromatin architecture in FGSCs, we performed *in situ* Hi-C^23^ with two biological replicates of FGSCs and other cells (SSCs, NSCs, iPSCs and STOs) generating approximately 400 million reads for each replicate. After filtering artificial reads and normalization, we obtained a total of over 2 billion valid Hi-C reads, including an average of 1 billion long-range (>20 kb) intra-chromosomal *cis* contacts and 400 million inter-chromosomal *trans* contacts (Table S1). We confirmed the high reproducibility of the Hi-C data (Figure S1) and combined the two biological replicates into a single set of merged Hi-C data per cell type, to reach a maximum resolution of 20 kb.

Analysis of the Hi-C data showed that the high-order chromatin organization of the whole genome in FGSCs differed from that in the other cells (Figure S2). An overview of the intra-chromosomal contact heat maps revealed that FGSCs displayed a distinct chromatin organization (Figure 2A). We further examined the characteristics of the chromatin organization by analyzing the patterns of compartment status and TADs in the autosomes across cells, avoiding sex chromosome effects. The compartment status was classified as active (A) or inactive (B) (Table S2). FGSCs were more similar to SSCs and NSCs in terms of A/B compartments, compared with iPSCs and STOs (Figure 2B), suggesting that FGSCs were ASCs. The patterns of TADs and directional indexes (DI) were almost the same for these cells (Figure 2B). We counted the numbers of compartments and TADs in the cells and showed that FGSCs had the lowest number of TADs, and the number of compartments was similar to that in SSCs (Figure 2C and Table S3). We also calculated the average intra-chromosomal contact probability of cells and found that the chromatin interaction frequency decreased monotonically from 10^5^ to 10^8^ bp for FGSCs and the other cells (Figure 2D). The contact probability curves were similar in five cell types from 10^5^ to 10^6.5^ bp, but changes occurred at long-distance genome, as reported previously^18^.

**Figure 2.**
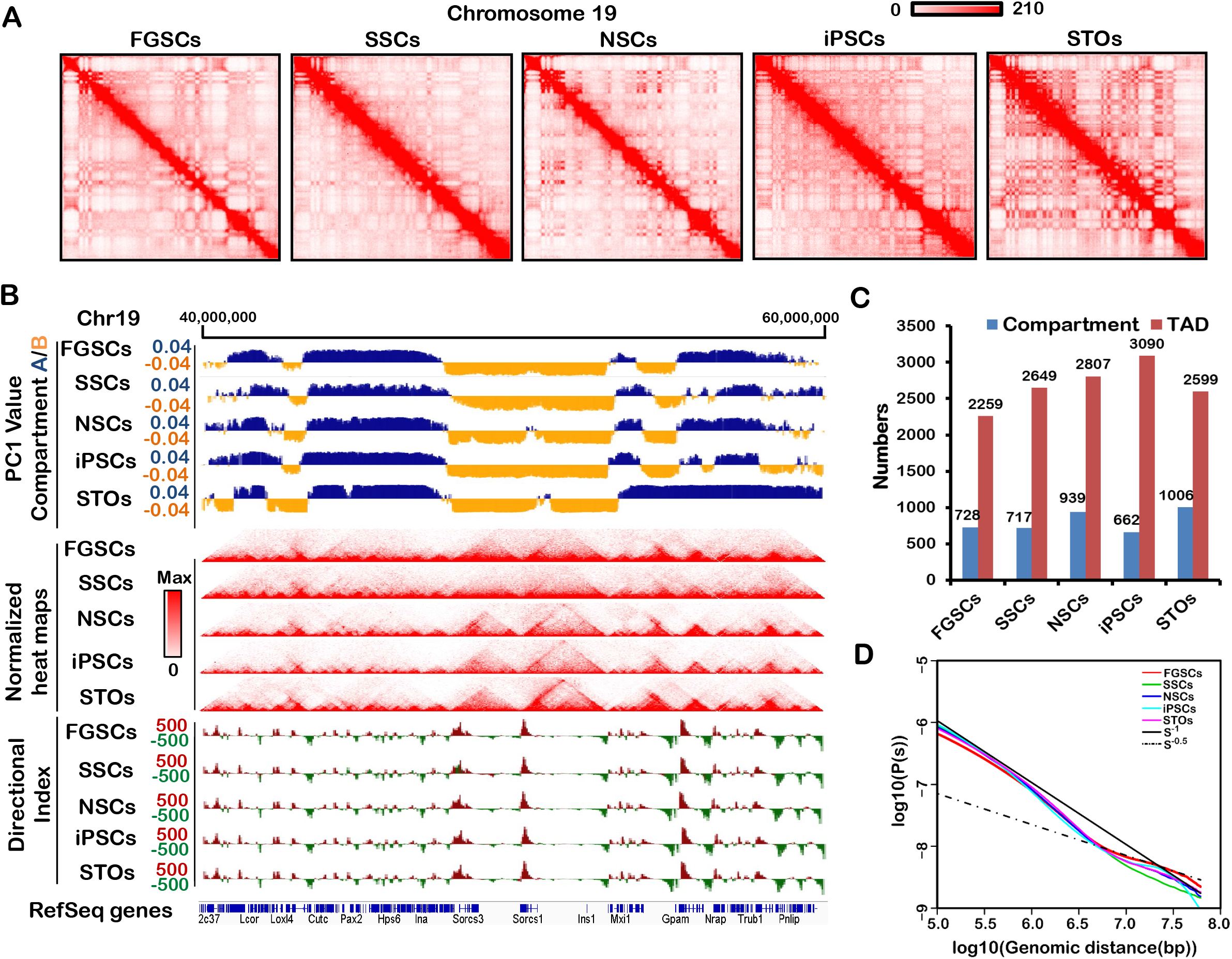
Overall chromosome structure in FGSCs. (A) Contact matrices from chromosome 19 in FGSCs, SSCs, NSCs, iPSCs and STOs. (B) First principal component (PC1) value, normalized Hi-C interaction heat maps, and directional indexes (DIs) in FGSCs and other cells at 20-kb resolution. PC1 value was used to indicate A/B compartment status, where a positive PC1 value represented the A compartment (blue) and a negative one represented the B compartment (yellow). (C) Numbers of identified A/B compartments and TADs in FGSCs and other cells. (D) The average contact probability across the genome decreased as a function of genomic distance.

### FGSCs Exhibited Distinct Compartment Status

Systematic analysis of the compartment status in FGSCs showed that genes had higher expression levels in compartment A than in compartment B (Figure S3A), indicating that compartment status was correlated with gene expression. Combining with the ChIP-Seq data analysis, we observed that H3K27ac and H3K4me3 were highly correlated with compartment status (Figure S3B). The genome browser showed that H3K27ac was highly enriched in compartment A but not in compartment B (Figure S3C). By k-means clustering of the compartment status in FGSCs compared with other cells, we obtained activate and inactive compartments of FGSCs (Figure 3A). Furthermore, switching compartments of FGSCs accounted for a high proportion (about 50%) of the total number in the genome compared with iPSCs and STOs, but a smaller proportion (about 30%–40%) compared with SSCs and NSCs (Figure S4A). These results suggested that FGSCs had a unique A/B compartment status that was more similar to other ASCs than to iPSCs or STOs. Additional RNA-Seq data revealed that the genes located in the switching compartment tended to be significantly differentially expressed compared with the stable compartments (Figure 3B). This was consistent with ChIP-Seq signal results, which showed dramatic differences of H3K27ac and H3K4me3 enrichment in the switching compartment compared with the stable compartment (Figure S4B). By calculating the covariation between gene expression and compartment status, we identified a subset of 1206 genes that were highly correlated with compartment status (Table S4). Gene Ontology (GO) analysis showed that these genes were particularly associated with stem cell population maintenance and cell proliferation (Figure S4C). Among these, *Coprs*, as an ASC marker located in the A compartment of FGSCs, SSCs, and NSCs, showed higher expression than in iPSCs in which it was located in the B compartment, consistent with a previous report^24^ (Figure 3C). In addition, *Nanog* as a pluripotent stem cell marker, showed higher expression in iPSCs, being located in the A compartment in iPSCs while switching to the B compartment in other types of cells (Figure 3C). These findings suggested that FGSCs had a unique compartment status characteristic of ASCs, which could work together with histone modification to regulate gene expression to determine their features.

**Figure 3.**
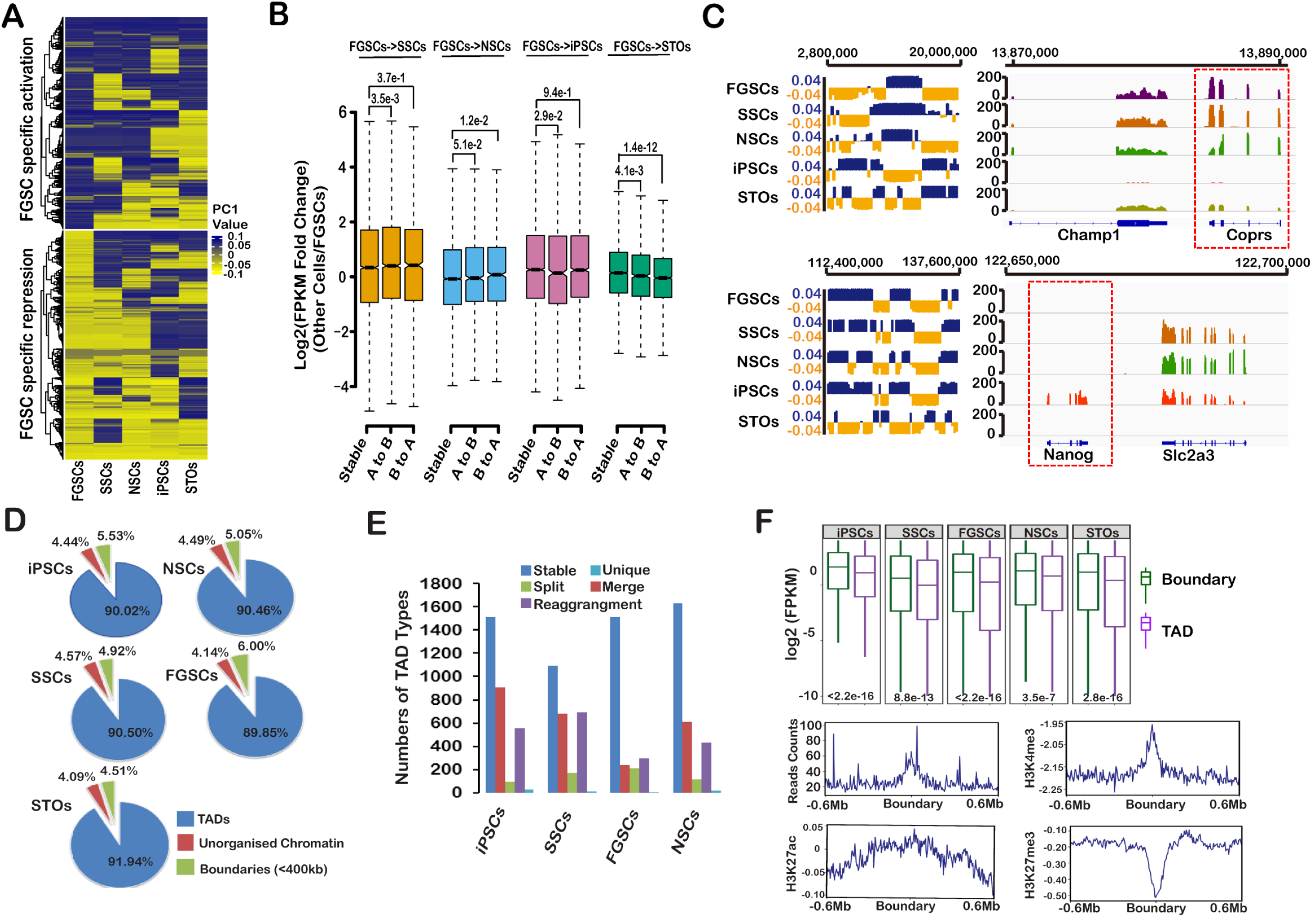
FGSCs exhibited specific compartment status. (A) K-means clustering (k=2) of PC1 values of the genome that change A/B compartment status in FGSCs. (B) Genes that changed compartment status (A to B or B to A) or that remained the same (stable) compared with FGSCs (P value by Wilcoxon’s test). (C) IGV snapshots for *Coprs* and *Nanog* showed concordance between their expression and PC1 values. (D) Percentages of TADs and TAD boundaries in the genome. (E) Numbers of TAD types, including stable, merged, split, unique, and rearrangement, in stem cells. (F) Comparison of gene expression between TADs and TAD boundaries. Genome-wide average distribution of RNA-Seq, H3K27ac, H3K4me3, and H3K27me3 reads around the domain boundaries in FGSCs.

We then identified the TADs in FGSCs using DI (Table S3). Well-defined TADs were conserved in ~90% of the genome across cell types (Figure 3D). We classified the TADs into five types: stable, merge, split, rearrangement, and unique. Most of the TADs belonged to the stable type (Figure 3E), suggesting that TAD structure was highly stable across all five types of cells, in accordance with a previous report^25^. However, the absolute DI showed that the strength of the TADs differed between FGSCs and the other cells (Figure S5A), suggesting that, although TAD domains were stable, FGSCs had a distinct frequency of intra-TAD interactions. We further explored the relationship between TADs and gene expression, and observed that gene expression was higher in TAD boundaries than in TADs in FGSCs and other cells (Figure 3F), illustrating that genes were more activated in these TAD boundaries (Figure S5B). We classified the boundaries into cell-type-specific and common boundaries, and identified 417, 369, 286, 263, and 48 cell-type-specific boundaries and 833 common boundaries, suggesting that most boundaries were stable across cell types. Further study of the relationship between cell-type-specific boundaries and gene expression indicated that gene expression changed significantly between specific and common boundaries (Figure S5C). These results revealed that FGSCs had stable TADs, but that gene expression was activated in the TAD boundary.

### Cell-type-specific Chromatin Loops Revealed FGSC Signature

We systematically analyzed the chromatin loops and identified 4832, 1906, 7004, 3060, and 6951 chromatin loops in FGSCs, SSCs, NSCs, iPSCs and STOs, respectively. Using a Venn diagram, we observed that only a few (n=177) chromatin loops were shared across all cell types (cell-type shared loops) (Figure 4A), suggesting that most chromatin loops were cell-type-specific and that FGSCs had distinct chromatin loops. Furthermore, genes located in these cell-type-specific loops were highly enriched in cell-type-related GO categories (Figure 4B). For instance, *Sox17*, as a transcription factor involved in embryonic development^26^, formed chromatin loops in iPSCs but gradually disappeared in FGSCs, SSCs, NSCs, and STOs, which was also most highly expressed in iPSCs (Figure S6A), suggesting that its expression could be regulated by *cis* regulation of the chromatin loops. Based on previous results, both A/B compartment status and TADs could affect gene expression^27,28^. We therefore investigated if the formation of cell-type-specific chromatin loops was related to compartment A/B status or to TADs, and found that 60% of cell-type-specific loops were commonly localized in the switching compartments (Figure S6B), while about 30% were localized in the stable TAD type (Figure S6C). This suggested that the chromatin loops relied more on compartment switching to regulate gene expression, and were not dependent on TAD type to exert their function.

**Figure 4.**
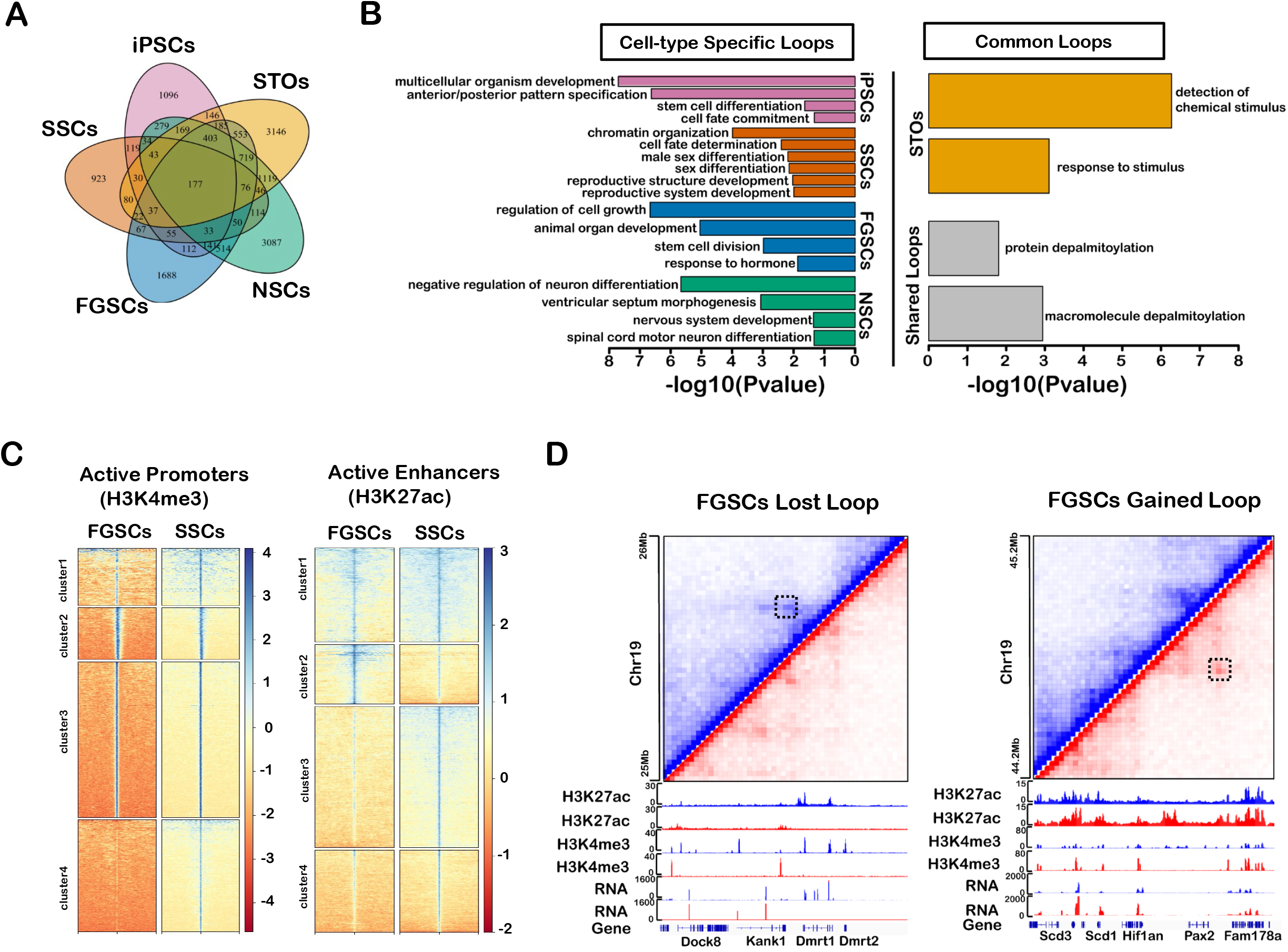
Cell-type-specifîc chromatin loops revealed FGSCs signature. (A) Venn diagram showing that most chromatin loops were cell-type-specific in five types of cells. (B) GO enrichment of cell-type-specific chromatin loops. (C) K-means clustering of H3K4me3 and H3K27ac ChIP-Seq data at promoter and enhancer regions between FGSCs and SSCs. (D) Chromatin loops gained or lost in FGSCs compared with SSCs.

We further examined the specific chromatin loops in FGSCs as GSCs by comparing them with the loops in SSCs using ChIP-Seq data. K-means clustering of ChIP-Seq signals identified four major classes on active promoter (H3K4me3) and enhancer (H3K27ac) sites (Figure 4C). Cluster 4 of active promoters and most of the active enhancers were obviously different between FGSCs and SSCs (Figure 4C).

Meanwhile, GO enrichment analysis further revealed that specific loops with different active promoters and enhancers were enriched in reproductive processes such as sex differentiation, sexual reproduction, and reproductive development (Fisher’s exact test, P < 0.05; Figure S7). We divided these loops into two subsets: FGSCs lost and FGSCs gained (Figure 4D). *Dmrt1*, as a conserved transcriptional regulator in male germ cells required for the maintenance and replenishment of SSCs^29^, was looped in SSCs but not in FGSCs. *Hif1an*, as a heterodimeric transcription factor related to interface with stem cell signaling pathways^30^, gained loops in FGSCs and could be related to female germline cell development (Figure 4D). Overall, these findings indicated that FGSCs possessed cell-type-specific chromatin loops that provided the spatial space for histone modification to regulate gene expression.

### X-Chromosome Structure Was Similar Between SSCs and FGSCs

One female X chromosomes is randomly inactivated during mammalian development to ensure matched dosages in males and females^31^. To dissect the high-order organization of the X chromosome in SSCs and FGSCs, we performed Pearson’s correlation analysis of the Hi-C matrix in the X chromosome. The results showed that SSCs and FGSCs had a strong correlation (Figure 5A). Furthermore, upon extracting the eigenvectors of the whole chromosome interactions and using the first principal component (PC1) score to compare the structure of the X chromosome in SSCs and FGSCs, we found that the X chromosome was more similar (correlation = 0.87) than the autosomes (mean correlation = 0.21) between SSCs and FGSCs (Figure 5B and Figure S8). By analyzing the PC1 score of the X chromosome in FGSCs, SSCs, and female embryonic stem cells (fESCs), considering that fESCs have two activated X chromosomes (Xa), we observed that FGSCs were more highly correlated with SSCs than with fESCs in the X chromosome (Figure 5C). This suggested that the X chromosome is similar between SSCs and FGSCs, probably due to one of the X chromosomes being inactivated (Xi) in FGSCs^32,33^.

**Figure 5.**
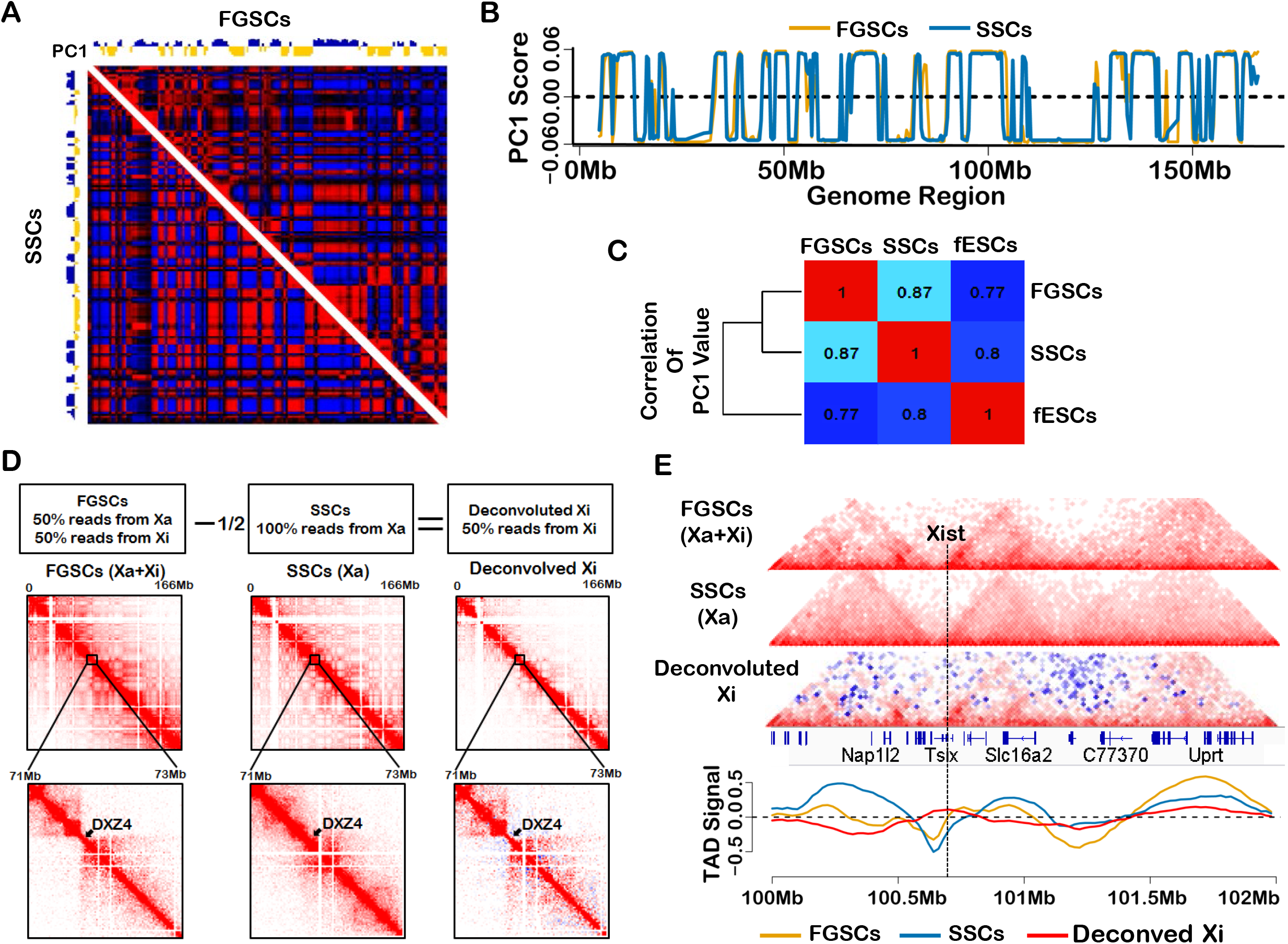
X-Chromosome structure conformation between SSCs and FGSCs. (A) Pearson’s correlation heat map showing similarity between FGSCs and SSCs. (B) PC1 values in the X chromosome in FGSCs and SSCs. (C) Spearman’s correlation of PC1 values in the X chromosome in FGSCs, SSCs, and fESCs. (D) Deconvolution of Xi signal from Hi-C data obtained in FGSCs (Xa+Xi) by subtracting the Xa contribution estimated from SSCs. A small region containing *Dxz4* showed two domains in FGSCs and deconvoluted Xi, but not in SSCs (blue pixels represent negative values, red pixels represent positive values). (E) Normalized chromatin interaction maps around *Xist* at 20-kb resolution. Plots show TAD signal (insulation score).

To identify the difference between the active X chromosome and the inactive one, we deconvoluted the respective Hi-C data of these chromosomes from FGSCs (Figure 5D). As expected, we visualized that the X chromosome was separated by a region containing the DXZ4 macrosatellite (which reportedly plays a crucial role in shaping Xi-chromosome structure) into two parts in FGSCs^34^, as well as in deconvoluted Xi, whereas this was not the case in SSCs (Figure 5D). This demonstrated that one of the X chromosomes in FGSCs was inactivated. Deconvoluted Xi displayed that the long-range contacts were attenuated for intra-TADs and inter-TADs (Figure 5D), consistent with a previous report^32,33^. We next investigated the structure of the region containing Xist, which is a key factor for inactivation of the X chromosome. Notably, FGSCs were similar to SSCs in the Xist region, while in deconvolved Xi it was shown that the Xist region lost most long-range contacts and retained a sub-TAD-like structure (Figure 5E), in support of a previous report^32^ Taken together, our data suggested that one of the X chromosomes was inactivated in FGSCs, to maintain relative consistency with SSCs to balance the gene expression between males and females.

### TADs Were Attenuated and then Disappeared During Female Germline Cell Development

FGSCs are derived from primordial germ cells and undergo meiosis into GV oocytes, and then to MII oocytes. To explore the dynamic changes of TADs during female germline cell development, we applied Hi-C to obtain the data of whole genome of chromatin architecture in mouse GV oocytes (Figure S9). After analysis of Hi-C data for MII oocytes, zygotes, two-cell embryos, and eight-cell embryos, we observed that TADs were attenuated in GV oocytes, disappeared in MII oocytes, and recovered in two-cell embryos (Figure 6A). A snapshot of TAD signals also showed that the TAD strength was weakened during female germline cell development and reprogramed at early embryonic development (Figure 6B), in contrast to changes in TADs during spermatogenesis^35^.

**Figure 6.**
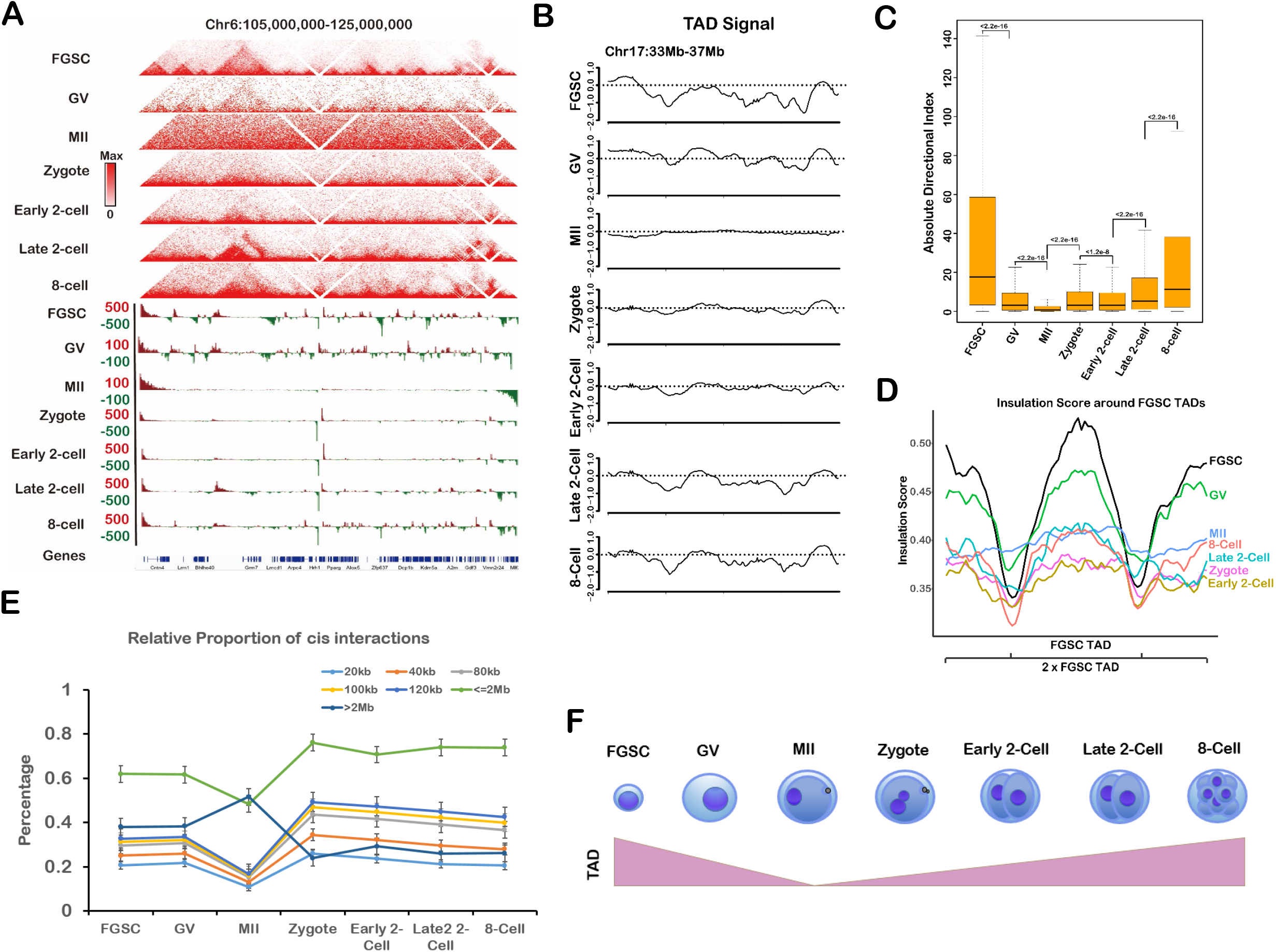
TADs were attenuated and then disappeared during female germline cell development. (A) Normalized Hi-C interaction frequencies during female germline cell development displayed as a heatmap. (B) TAD signals at chromosome 17:33–37 Mb during female germline cell development. (C) Boxplot of absolute DI values during female germline cell development. P value calculated by Kruskal-Wallis test. (D) Average insulation scores (IS) of different stages in female germline cell development at TADs (defined in FGSCs) and nearby regions. (E) Relative proportions of *cis* interactions at different genome distances versus total paired loci. (F) Graphical model for dynamic changes in TADs during female germline cell development.

We further examined the changes in TADs during female germline cell development by calculating the DI value, which reflected the degree of interactions of a given bin in upstream or downstream regions and was associated with calling TADs. Expectedly, the DI value was significantly reduced during female germline cell development and reestablished in early embryonic development (Figure 6C). Meanwhile, the insulation score showed that FGSCs had the strongest TAD boundaries, while these were decreased in GV oocytes and weakest in MII oocytes, and gradually increased in early embryonic development (Figure 6D). We further calculated the proportion of *cis*-short interactions (<2 Mb) and *cis*-long interactions (>2 Mb) versus total *cis*-interactions. The relative proportions of *cis*-short interactions in FGSCs was similar to GV oocytes, lowest in MII oocytes, highest in zygotes, and then reduced during early embryonic development, with similar interactions in eight-cell embryos and FGSCs (Figure 6E). Overall, these results demonstrated that TADs were attenuated and then disappeared during female germ cell development, and reestablished during early embryonic development (Figure 6F), thus revealing the pattern of TADs throughout female germ cell and early embryonic development.

### Identification of Conserved Allelic Chromatin Structures

Previous findings have reported the chromatin structure of the maternal genome is different with the paternal genome during early embryonic development^18,19^ We aimed to investigate how the conserved structures in these respective genomes affected their genome organization. Early embryonic Hi-C data were analyzed with single nucleotide polymorphisms between two mouse strains to track the maternal and paternal genomes. The correlation of compartment status according to the PC1 score, showed that the paternal genome was clustered in pachytene spermatocytes (PACs), sperm, paternal zygotes, and eight-cell embryos, while the maternal genome was clustered within MII oocytes, maternal zygotes, and eight-cell embryos (Figure 7A), indicating that the allelic chromatin structure was conserved in early embryonic development. FGSCs and SSCs, as early-stage germ cells, were clustered together (Figure 7A). Interestingly, the paternal genome clustered with the maternal genome at the two-cell embryo stage, but was completely separate at the eight-cell stage, suggesting that the two-cell stage plays an important role in allelic chromatin structure development, possibly because of the reestablishment of TADs at this stage. Furthermore, we observed some regions of compartments in allelic genome were conserved during the early embryonic development (Figure 7B). We then systematically identified the conserved A/B compartment regions by calculating Pearson’s correlation for the whole genome, with a sliding window of 2 Mb. The paternal genome had more conserved regions than the maternal genome (Figure 7C), indicating that the paternal genome was more conserved. Comparing the conserved regions, some allelic-specific conserved regions were identified by Venn diagrams (Figure 7D). By enriching for imprinted genes with Fisher’s exact test (P < 0.05), we found that the allelic-specific regions were significantly related to imprinted genes (Table S5) such as *Igf2r, Dlk1*, and *Dio3*, which were reported to highly express in maternal or paternal^36,37^ The results suggested that those imprinted genes were affected by the allelic-specific conserved region, which could regulate paternal or maternal development.

**Figure 7.**
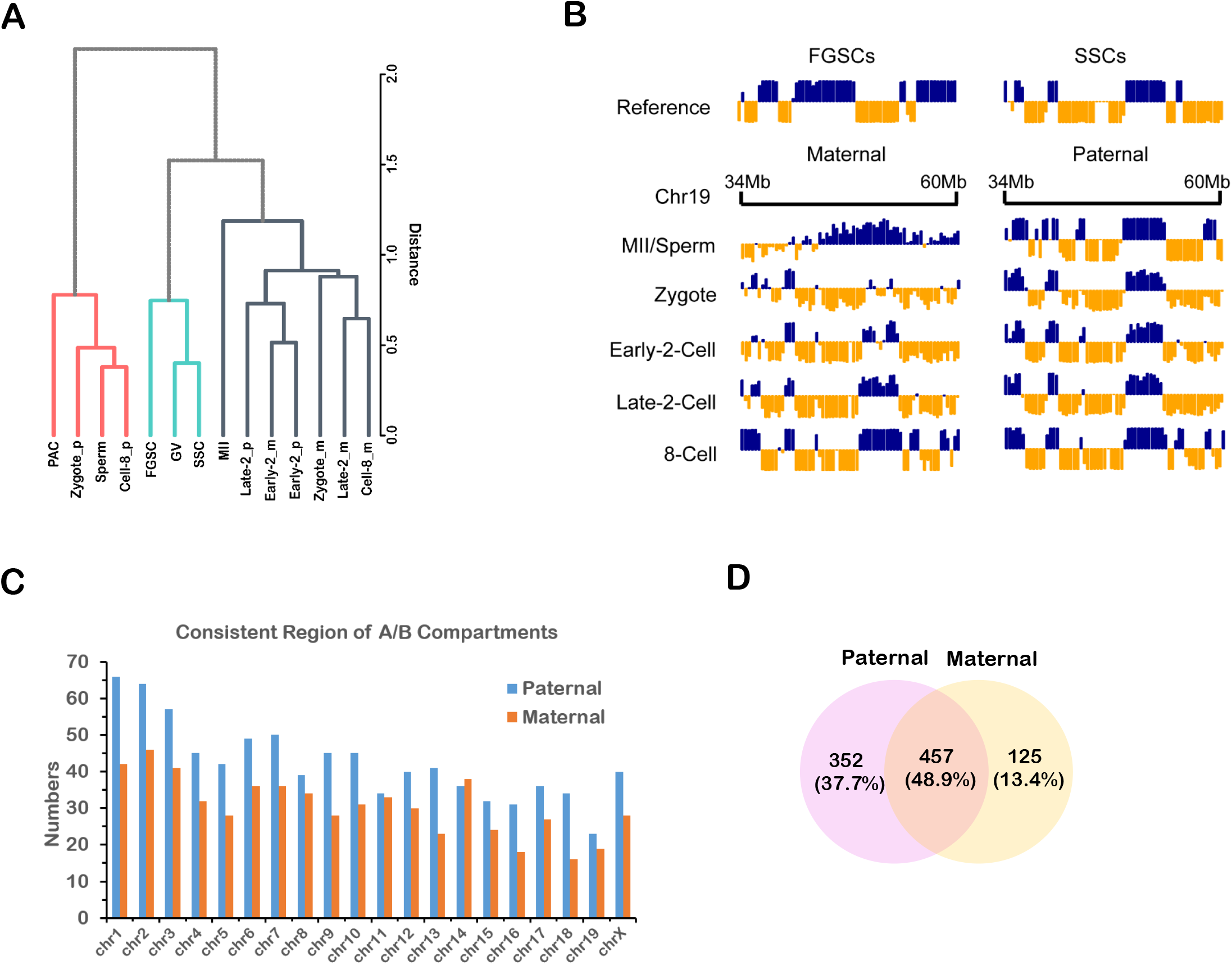
Identification of conserved allelic chromatin structures. (A) Hierarchical clustering of PC1 values based on maternal (black) and paternal (red) genome architectures. Reference shown in green. (B) Example of conserved compartment region at chromosome 19:34–60 Mb during early embryonic development. (C) Numbers of conserved compartment regions in paternal and maternal genomes. (D) Venn diagram showing overlap of conserved compartment regions between paternal and maternal genomes.

## DISCUSSION

Stem cells, including pluripotent stem cells and ASCs, have important implications in basic biology and regenerative medicine. However, regenerative medicine requires stem cells to be transplanted safely, with a particular focus on avoiding the development of cancer. Although gene mutations have been reported to be responsible for many diseases, including cancer^38,39^, recent studies have revealed that diseases such as cancer can also be caused by disruption of chromatin organization. The chromatin architecture thus plays crucial roles in preventing DNA damage, in gene mutation, and in ensuring appropriate gene transcription, DNA duplication, and developmental processes^40–42^. It is therefore essential that stem cells are characterized or identified in terms of their chromatin organization before their use in a clinical context. FGSCs not only have the abilities of self-renewal and differentiation, but are also responsible for passing on genetic information to the next generation. The stability of that genetic information is affected by the high-order genome organization^43^.

To identify the chromosome structure character of FGSCs, we compared FGSCs with pluripotent stem cells (iPSCs), ASCs (SSCs and NSCs) and somatic cells (STOs) by Hi-C technology. The results revealed that FGSCs had a distinct high-order genome structure in terms of the A/B compartment status, chromatin loops, and TADs. For further characterization, we identified FGSCs specific activated and repressed compartment regions, and obtained partially genes highly related with the switch of FGSCs compartment status. These genes were related to stem cell maintenance and differentiation pathways, strongly supporting the role of FGSCs as stem cells, with some shared characteristics with SSCs and NSCs, and belonging to ASCs. Moreover, FGSC-specific loops, which included active promoters and enhancers, could be significantly enriched in reproductive-related pathways. Among these, *hif1an*, which is related to Notch signaling, could be a potential marker for female germline cells. Our findings indicated that FGSCs belong to ASCs in the high-order organization, and further confirmed the existence of FGSCs, consistent with the previous reports for cellular and molecular characteristic^4–7,44,45^.

Previous studies indicated that both FGSCs and SSCs have their own unique epigenetic signatures^44,46^. We analyzed and compared the 3D genomic architectures of FGSCs and SSCs, and revealed that GSCs had their own unique high-order chromatin organization, especially in terms of A/B compartments and chromatin loops. Although FGSCs have one more X chromosome than SSCs, the architecture of the X chromosome in FGCSs was more similar to SSCs than to autosomes, suggesting that FGSCs maintain a balance of gene expression with SSCs by inactivating one X chromosome. Furthermore, analysis of the X-chromosome matrix of FGSCs indicated that it was separated into two domains by a region containing *Dxz4*, which has been reported to be an essential regulator of X chromosome inactivation^32^. Meanwhile, deconvoluted Xi data showed that *Xist* was also located in a region showing moderate interactions with a TAD-like structure, consistent with previous findings^32^. These results indicate that these differences could reflect Xi, supporting inactivation of one X chromosome in FGSCs^47^

Interestingly, recent studies reported that the chromatin architecture changed dynamically during spermatogenesis, with dissolved and then reappeared TADs and compartments^16,17^ On the contrary, TADs were attenuated and then disappeared in oogenesis. During early embryonic development, TADs recovered and started to appear in two-cell embryos, suggesting that this stage could play an essential role in genome organization, including paternal/maternal chromatin reprogramming. We therefore further analyzed paternal and maternal chromatin structures during early embryonic development, and found that the paternal genome clustered with the maternal genome at the two-cell stage, but was completely separate at the eight-cell stage, consistent with the above results.

In conclusion, we present a comprehensive overview of the chromatin organization of FGSCs to create a rich resource of high-resolution genome-wide maps. Our findings revealed that the chromatin architecture of FGSCs included unique features, especially in terms of compartment status and chromatin loops, which may contribute to their cell-type-specific gene regulation. These data will provide a valuable resource for future studies of the features of chromatin organization in mammalian stem cells, with important implications for their role in medical research and their potential and actual clinical applications.

## EXPERIMENTAL PROCEDURES

Additional information and details regarding this work may be found in the Supplemental Experimental Procedures.

## ACCESSION NUMBERS

The accession number for the expression and sequencing data reported in this paper is GEO: GSE126014 and GEO: GSE137771.

## SUPPLEMENTAL INFORMATION

Supplemental Information includes Supplemental Experimental Procedures, eight figures and four tables and can be found with this article online.

## AUTHOR CONTRIBUTIONS

T.G.G performed the Hi-C experiments, analyzed the data and wrote the manuscript. Z.X carried out ChIP-Seq experiments. X.W and W.L performed the NSC culture and identification. L.X undertook the FGSC culture and identification. W.Y performed the SSC culture and identification. L.H finished RNA-Seq experiment. H.C did the Hi-C experiment of GV stage oocyte. W.J and L.J supervised the experiment work and devised this study.

## ACKNOWLEDGMENTS

We thank Dr. Kang’s lab (Tongji University) for providing the iPSCs cell line. This work was supported by National Basic Research Program of China (2018YFC1003501, 2017YFA0504201), National Nature Science Foundation of China (81720108017, 81501316), the National Major Scientific Instruments and Equipment Development Project, National Nature Science Foundation of China (61827814).

