## Supplemental File for "Three-dimensional Genome Structure Reveals Distinct Chromatin Signatures of Mouse Female Germline Stem Cells During Development"

**Method**

**Animals**

The CAG-EGFP mice from the Model Animal Research Center of Nanjing University, also named C57BL/6-Tg (CAG-EGFP) C14-Y01-FM131-Osb ^1^. The Ddx4-Cre;mT/mG mice were generated as described previously ^2^. DBA/2 mice were obtained from the Shanghai Slac Laboratory Animal Co., Ltd. (Shanghai, China). All procedures involving animals were approved by the Institutional Animal Care and Use Committee (IACUC) at Shanghai Jiao Tong University, and all experiments were performed in accordance with the approved protocols.

**Isolation and Culture of SSCs**

Mouse SSCs were isolated and cultured from 6-day-old male F_1_ progeny of DBA/2 × CAG-EGFP mice, as previously described ^3^. In brief, SSCs (>20 passages) were cultured on mitomycin C (MMC)-treated mouse embryonic fibroblast (MEF) feeder cells with SSC culture medium. The SSC culture medium included StemPro-34 SFM supplemented with StemPro supplement (Invitrogen, Carlsbad, CA, USA), 100 μg/ml transferrin, 25 μg/ml insulin, 10 ng/ml human basic fibroblast growth factor (Invitrogen), 20 ng/ml recombinant human epidermal growth factor (Invitrogen), 10 ng/ml recombinant human glial cell line-derived neurotrophic factor (Invitrogen), 6 mg/ml D-(+)-glucose, 60 mMputrescine, 30 nM sodium selenite, 2 mM L-glutamine (Invitrogen), 30 μg/ml pyruvic acid, 1 μl/ml DL-lactic acid (Sigma, St. Louis, MO, USA), 5 mg/ml bovine serum albumin (Sigma), 10 μM 2-mercaptoethanol (Sigma), 1×MEM vitamin solution (Invitrogen), 1× nonessential amino acid solution (Invitrogen), 0.1 mM ascorbic acid, 10 μg/ml D-biotin (Sigma), 1% fetal bovine serum (Gibco), and 1× penicillin/streptomycin solution (Invitrogen). The medium was changed every 2–3 days.

**Isolation and Culture of FGSCs**

Mouse FGSCs were isolated and cultured from neonatal Ddx4-Cre;mT/mG mouse ovaries (6 days old), as previously described ^2^. Briefly, FGSCs (>18 passages) were cultured on mitotically inactivated STO (SIM mouse embryo-derived thioguanine- and ouabain-resistant) cell feeders in minimum essential medium alpha (MEMα; Invitrogen) supplemented with 10% fetal bovine serum (FBS) (Life Technologies), 10 ng/mL mouse leukemia inhibitory factor (Santa Cruz Biotechnology), 20 ng/mL mouse epidermal growth factor (EGF) (PeproTech), 10 ng/mL basic fibroblast growth factor (bFGF) (PeproTech), 10 ng/mL mouse glial cell line-derived neurotrophic factor (GDNF) (PeproTech), 1 mM nonessential amino acids (Life Technologies), 2 mM L-glutamine (Sigma), 30 mg/mL pyruvate (Amresco), and 50 mMβ-mercaptoethanol (Biotech). The FGSCs were subcultured every 4–7 days.

**Isolation and Culture of NSCs**

NSCs were isolated from E12.5 mouse embryonic cortex, in accordance with a previously described procedure ^4^^,^^5^. The primary NSCs were seeded onto poly-L-ornithine (Sigma-Aldrich)- and laminin (Invitrogen)-coated dishes, cultured as monocultures. Neural basal medium with 20 ng/ml EGF (PeproTech), 20 ng/ml bFGF (PeproTech), 20 ng/ml heparin (Sigma-Aldrich), and 2% B27 (Invitrogen) was used as NSC proliferation medium. For differentiation, NSCs were seeded on poly-L-ornithine (Sigma-Aldrich)- and laminin (Invitrogen)-coated dishes. Upon NSC attachment, the medium was changed to differentiation medium identical to the NSC proliferation medium without growth factors (EGF and bFGF). After differentiation for 10 days, neural- and glial-specific markers were used to determine the differentiation potential of cultured NSCs.

**STO Culture**

STOs were maintained in Dulbecco’s modified Eagle’s medium (DMEM) with high glucose (Life Technologies), 10% FBS (Life Technologies), 1% nonessential amino acids (Life Technologies), 2 mM glutamine (Sigma), and penicillin (100 U/ml; Sigma)/streptomycin (0.1 mg/ml; Sigma) at 37°C and 5% CO_2_, and passaged after 3–4 days.

**Immunofluorescence**

The immunofluorescence procedure was performed as described previously with minor modification ^6^. Cultured cells were washed twice with phosphate-buffered saline (PBS) and then fixed in 4% paraformaldehyde for 15 min. After that, cells were rinsed twice and incubated with PBS containing 0.1% (v/v) Triton X-100 for 15 min. Fixed cells were blocked by 10% normal horse serum in PBS for 10 min at room temperature. The cells were then incubated with primary antibodies: mouse monoclonal anti-NESTIN and rabbit polyclonal anti-SOX2 (1:180 dilution; Abcam), mouse polyclonal anti-TUJ-1 and rabbit polyclonal anti-GFAP (1:150 dilution; Abcam), mouse monoclonal anti-MVH (1:500 dilution; Abcam), and mouse monoclonal anti-PLZF (1:100 dilution; Santa Cruz Biotechnology). After 1 h of incubation at room temperature with the primary antibodies, the cells were rinsed twice in PBS and then incubated in the dark with a fluorescein isothiocyanate-conjugated secondary antibody, either goat anti-rabbit IgG or goat anti-mouse IgG (1:200 dilution; Proteintech, Chicago, IL, USA) for 60 min at 37°C. This was followed by rinsing and staining of the nucleus with 4′,6-diamidino-2-phenylindole (DAPI) (1:1000 dilution; Sigma)-containing PBS for 10 min at room temperature. After washing twice with PBS, the cells were examined in fresh PBS under an inverted fluorescence microscope (Leica).

**RT-PCR**

Total RNA from cells was extracted using Trizol reagent (Invitrogen), in accordance with the manufacturer’s instructions. After extraction, 1 µg of total RNA was used to synthesize cDNA with Primescript Reverse Transcriptase (ThermoFisher). Specific marker genes, such as *Nestin*, *Sox2*, *Pax6*, *Olig2*, *Ascl1*, *Exv5*, *Plzf*, *Mvh*, *Oct4*, *Fragilis*, *Stella*, *Dazl*, and *Gfra1*, were used to characterize the cells. *Gapdh* was used as an internal control. Primer sequences are listed in Supplementary Table S6.

**Mouse** **Germinal Vesicle Oocyte Collection**

Six-week-old C57BL/6J female mice were intraperitoneally injected with 10 IU pregnant mare serum gonadotropin and then killed by cervical dislocation after 46-48h. Mouse germinal vesicle (GV) oocytes were isolated by puncturing of ovaries with hypodermic needles. To avoid the granulosa contamination, the GV oocytes were washed eight times with MEM medium containing 1 mg/mL hyaluronidase. The zona pellucida was then removed usingAcidic Tyrode's solution (Sigma) followed by fixing in formaldehyde ^7^. GV oocytes were then prepared for Hi-C experiments.

***In Situ* Hi-C Library Generation**

Cells were used for *in situ* Hi-C, including iPSCs (from Kang’s Lab, Tongji University), FGSCs, SSCs, NSCs, and STOs ^8^. Hi-C libraries were generated in accordance with the standard *in situ* Hi-C protocol, with minor modification ^9^. Five million cells were harvested with 0.05% trypsin, washed twice, resuspended with 22.5 ml of DMEM medium, and then crosslinked with 62.5 µl of 37% formaldehyde (F8775; Sigma) to a final concentration of 1% for 10 min at room temperature. Formaldehyde was quenched by adding 1.25 ml of glycine to a final concentration of 0.2 M and the cells were incubated for 5 min at room temperature, then transferred to ice for 20 min. The fixed cells were centrifuged at 400×g and 4°C and washed with cold PBS once, followed by storage at −80°C. Fixed cells were resuspended in 0.5 ml of lysis buffer and then incubated on ice for 30 min. Nuclei were pelleted by centrifugation at 3000×g for 5 min and washed once with 500 μl of cold lysis buffer. The pellet was resuspended in 50 μl of 0.5% sodium dodecyl sulfate (SDS) and incubated at 62°C for 5–10 min. Then, SDS was quenched by adding 145 μl of water and 25 μl of 10% Triton X-100 and incubated at 37°C for 15 min. The nuclei were digested overnight at 37°C with 20 μl of Mbol restriction enzyme (NEB, R0417) and 25 μl of 10× NEBuffer2. The next day, the nuclei were incubated at 62°C for 20 min, followed by cooling to room temperature. A mixed solution (50 μl) (37.5 μl of 0.4 mM biotin-14-dATP, 1.5 μl of 10 mM dCTP, 1.5 μl of 10 mM dGTP, 1.5 μl of 10 mM dTTP, and 8 μl of 5 U/μl DNA Polymerase I) was then added and incubated at 37°C for 2 h. Ligation master mix (663 μl of water, 120 μl of 10× NEB T4 DNA ligase buffer, 100 μl of 10% Triton X-100, 12 μl of 10 mg/ml bovine serum albumin, and 5 μl of 400 U/μl T4 DNA ligase) was added to a volume of 900 μl and slowly rotated for 6 h at room temperature. Next, to degrade protein, 50 μl of 20 mg/ml proteinase K and 120 μl of 10% SDS were added and incubated at 55°C for 30 min. Then, 5 M sodium chloride (130 μl) was added and incubated at 68°C overnight. On the third day, the DNA was purified by adding 1.6× volumes of pure ethanol and 0.1× volumes of 3 M sodium acetate, pH 5.2. Subsequently, DNA was diluted with 500 μl of Tris buffer and sheared by a Digital Sonifier (Branson). Dynabeads MyOne Streptavidin T1 beads (50 μl) (Thermo Scientific) were added and washed once with 400 μl of 1× Tween Washing Buffer (TWB) [5 mM Tri-HCl (pH 7.5), 0.5 mM EDTA, 1 MNaCl, 0.05 Tween 20), separated on a magnet, and then the solution was discarded. The beads were resuspended in 300 μl of 2× Binding Buffer [10 mMTris-HCl (pH 7.5), 1 mM EDTA, 2 MNaCl], mixed with the DNA, and incubated at room temperature for 1 h with rotation, followed by washing twice with 500 μl of TWB and reclaiming the beads using a magnet. The beads were next resuspended with a mixture [88 μl of 1× NEB T4 DNA ligase buffer with 10 mM ATP, 2 μl of 25 mM dNTP mix, 5 μl of 10 U/μl NEB T4 PNK, 4 μl of 3 U/μl NEB T4 DNA Polymerase I, 1 μl of 5 U/μl NEB DNA Polymerase I, Large (Klenow) Fragment] and incubated at room temperature for 30 min, followed by separation on a magnet and washing twice with 1× TWB. The beads were resuspended with 100 μl of mixture (90 μl of 1× NEBuffer 2, 5 μl of 10 mM dATP, 5 μl of 5 U/μl NEB KlenowExo Minus) and incubated at 37°C for 30 min. Then, separation on a magnet was performed, after which the solution was discarded and washing was carried out twice. The beads were resuspended in 50 μl of 1× NEB Quick ligation reaction buffer (B6058; NEB), and 2 μl of NEB DNA Quick ligase (M2200; NEB) and 3 μl of an Illumina indexed adapter of NEBNext Multiplex Oligos for Illumina (E73358; NEB) were added at room temperature for 15 min. Then, 3 μl of USER enzyme of NEBNext Multiplex Oligos for Illumina was added to the ligation mixture at 37°C for 15 min. The DNA was washed twice and eluted by resuspending the beads with 50 μl of 10 mMTris-HCL (pH 8.0), followed by incubation at 98°C for 10 min. The DNA suspension was transferred into a fresh tube and stored at −20°C. The Hi-C library was amplified in a 50-μl PCR system (25 μl of NEBNext Q5 Hot Start HiFi PCR mix, 5 μl of index primer of NEBNext Multiplex Oligos for Illumina, 5 μl of universal primer of NEBNext Multiplex Oligos for Illumina, and 15 μl of DNA template) with the following PCR conditions: initial denaturation at 98°C for 30 s; eight cycles of denaturation at 98°C for 10 s and annealing/extension at 65°C for 75 s; followed by final extension at 65°C for 5 min, and then maintenance at 4°C. The DNA fraction in the size range of 300–500 bp was selected using AgencourtAMPure XP beads (Beckman Coulter). Briefly, the PCR product was added into a total volume of 250 μl with 1× Tris-HCl, and 175 μl of AMPure XP beads were added to the PCR product (0.7 volumes). Mixing was then performed by pipetting, followed by incubation at room temperature for 5 min. The beads were washed once with 700 μl of 70% ethanol without mixing. They were then resuspended with 100 μl of 1× Tris-HCl buffer and another 70 μl of AMPure XP beads were added. Mixing was performed by pipetting, followed by incubation at room temperature for 5 min. The beads were then washed twice with 700 μl of 70% ethanol without mixing. They were subsequently left on the magnet for 5 min to allow the remaining ethanol to evaporate. Finally, the DNA was eluted with 33 μl of 1× Tris-HCl buffer and incubated at room temperature for 5 min, followed by separation on a magnet and transfer of the solution to a fresh labeled tube. This produced the final Hi-C library. The quality of the Hi-C library was determined using the QubitdsDNA HS Assay and Agilent 2100 DNA 1000 HS kit. The high-quality libraries were sequenced using an Illumina sequencing platform.

**ChIP-Seq Library Preparation**

The preparation of ChIP and input DNA libraries was performed as previously described^10^. In brief, two cells were crosslinked with 1% formaldehyde for 5 min at room temperature and quenched with 125 mM glycine. Cells were then put on ice, resuspended in cold cell lysis buffer [140 mM NaCl, 1 mM EDTA pH 8.0, 1% TritonX-100, 0.1% SDS, and protease inhibitors (Roche)]. Nuclei were sonicated into fragments of 200–1000 bp in size. The chromatin fragments were pre-cleared and then immunoprecipitated with Protein A+G magnetic beads coupled with anti-H3K4me3 (ab8580; Abcam), anti-H3K27ac (ab4729; Abcam), and anti-H3K27me3 (07-449; Millipore). After reverse crosslinking, immunoprecipitated DNA and input DNA were end-repaired and adapters were ligated to the DNA fragments using NEBNext Ultra End-Repair/dA-Tailing Module (E7442; NEB) and NEBNext Ultra Ligation Module (E7445; NEB). High-throughput sequencing of the ChIP fragments was performed using Illumina NextSeq 500, following the manufacturer’s protocol.

**Hi-C Data Processing, Mapping, and ICE Normalization**

Hi-C pair-end was trimmed of adaptor sequences and low-quality reads were filtered with BBmap (version 38.16). HiCPro (version 2.7) ^11^ was then used to map, process, and perform iterative correction for normalization. Reads were independently aligned to the mouse reference genome (mm9) with the bowtie2 algorithm^12^. Uncut DNA reads, re-ligation reads, continuous reads, and PCR artifacts were discarded. We then constructed a contact matrix using the unique mapped reads (MAPQ>10). We divided the genome into sequential bins of equal size and valid read pairs were then binned at a specific resolution. ICE ^13^normalization was applied to remove bias in the raw matrix, such as GC content, mappability, and effective fragment length in Hi-C data.contact matrices were finally generated at binning resolutions of 10, 20, 40, 200, and 400 kb.

**Validation of Hi-C Data**

Data reproducibility was confirmed by calculating Pearson’s correlation coefficient between the two Hi-C repeats. For each possible interaction I_ij_ between two replicates, these were correlated by comparing each point’s interaction in the normalized interaction matrix. Considering that the interaction matrix was highly skewed toward proximal interactions, we calculated the correlation to a maximum distance restricted to 2 Mb between points i and j. R was used to calculate Pearson’s correlation between two duplicates.

**Contact Probability *P(s)* Calculation**

*P(s)*only considering intra interactions, was calculated with normalized interaction matrices at 40-kb resolution, as described previously ^14^. Briefly, we divided the genome into 40-kb bins and counted the number of interactions at corresponding distances for each distance (separated by 40, 80, 120, 160 kb, etc.). We then divided the number of interactions in each bin by the total number of possible region reads as *P(s)*. Finally, we normalized the sum of *P(s)* over the range of distances as 1. The curve (log–log axis) was generated by locally weighted scatterplot smoothing.

**Identification of A and B Compartments**

The R package (HiTC) ^15^ was used to generate the PC1 eigenvectors using 400-kb normalized matrices with pca.hic function, usingthe options: (normPerExpected=TRUE, npc=1), for which a positive value indicates the A compartment and a negative value indicated the B compartment. To investigate compartment switching, we defined switched bins only if the PC1 eigenvectors changed in the same direction for two replicates.

**Identification of Concordant Genes with A/B Compartment Switch**

We defined genes with concordant changes in expression and compartment status according to a previously described method, with minor modifications ^16^. Briefly, the covariance between the vector of the gene expression values (FPKM) and the vector of PC1 values for each gene was calculated across five cell types. We then used the covariance metric to quantitatively define ‘concordance’. The observed covariance values were compared to a random background distribution to calculate a P value for the covariance for each gene. Randomly shuffling the vector of FPKM for each gene produced the background distribution, and then obtained the covariance between the PC1 values and the random gene expression vector. A rank-based P value could be calculated for the observed covariance values with 1000 repeats for each gene. Concordant genes were defined as those with a P value < 0.01.

**TAD Calling, TAD Boundaries, and TAD Types**

We calculatedthe location of the TADs using the directional index (DI) value, as previously described^17^. We first calculated the DI value for each bin based on the ICE-normalized and depth-normalized matrix and then used this value as the input for a Hidden Markov Model (HMM) to call TADs. TAD boundaries were defined as those <400 kb. To analyze TAD types, we calculated the proportion of overlapping TADs compared with STOs using bedtools (version 2.25.0) ^18^. Two or more TADs in a stage fused into one TAD in STOs wa defined as “merge”; one TAD in a stem cell divided into two or more TADs in STOs was defined as‘split’; and a TAD was unique to stem cells and not in STOs was defined as ‘unique’. In addition to the merge, split, and unique TADs, a proportion of overlapping TADs between stem cells and STOs > 0.70was defined this as ‘stable’, and other TADs were defined as ‘rearrangement’. We identified cell-type-specific boundaries in stem cells by comparing the boundaries between stem cells and STOs. Boundaries located within 10× bins (200 kb) were considered as ‘common’, while boundaries located at >200 kb were considered as ‘cell-type-specific’.

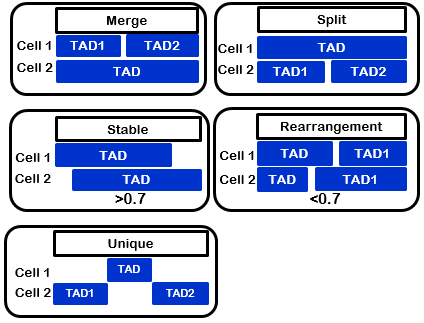

**TAD Signal Calculation**

We calculated TADs signals using the insulation score ^19^. The insulation score for each bin in the 20kb was calculated by the average number of interactions that occurred across each bin. Using this matrix, we then plotted the insulation score distribution centered in the FGSC TADs (up/down to 0.5 TAD).

**Identification of Enhancers and Promoters in FGSCs**

We carried out chromatin loop calling using the tool (PSYCHIC), as described previously ^20^, with the following parameters: -res 20,000 -win 2,000,000. Chromatin loops were defined when the P <0.0001 and the relative fold change (observation interactions/expected interactions) was >2. Active promoters and active enhancers were merged with ChIP-Seq data for H3K4me3 and H3K27ac, respectively.

**Calculation of Conserved Compartment Region**

Pearson’s correlation was used to identify conserved compartment regions. Briefly, using a PC1 score matrix at a resolution of 400kb, we slid 2-Mb windows along whole genome to calculate the Pearson’s correlation between two development stages and calculated the mean value of this correlation. Regions with R>0.6 were chosen as conserved compartment region during early development.

**ChIP-Seq Data Analysis**

We aligned fastq files to the mm9 reference genome, removed PCR duplicates using Samtools (version 2.0.1)^21^, and generated normalized genome coverage tracks from uniquely mapping reads (MAPQ>10) using deepTools2 (version 3.1)^22^. Biological replicates were pooled, and coverage was then calculated as the average reads per million mapped reads (RPM) in 1-kb bins. To identify the correlation between ChIP-Seq and A/B compartment, we summed the log2 of the fold enrichment (treatment/input) from ChIP-Seq to calculate the relative ChIP-Seq signal in each compartment.

**RNA-Seq Library Generation and Data Analysis**

Total RNA was extracted from 2–6 million cells using Trizol Reagent (Invitrogen). The RNA quality was assessed using Agilent Bioanalyzer 2100. RNA-Seq libraries were prepared using the KAPA Stranded mRNA-Seq kit, following the manufacturer’s instructions. After preparation, libraries were quantified using a Qubit fluorometer and sequenced with HiSeq Platform (2× 150 bp). All RNA-Seq data were trimmed and aligned to the mm9 reference genome using Hisat2 (version 4.8.2) ^23^with the default parameters. Gene expression FPKM was calculated by Cufflinks (version 2.2.1) ^24^using the RefSeq database from the UCSC genome browser. Sequencing depth was normalized.

**X-Chromosome Analysis**

To estimate the contribution of the inactive X chromosome (Xi) to the Hi-C signal obtained in FGSCs, we used a previously described method with minor modification ^25^. First, we normalized read numbers obtained using R package (limma) ^26^with quantile normalization in FGSCs and SSCs to accommodate differences in frequency strictly due to sequencing depth. We then divided the normalized SSC signal by two, to subtract only the activated X-chromosome (Xa) component from the FGSC (XaXi) data (and not 2× Xa). Assuming that half of the total Hi-C reads in FGSCs (XaXi) were obtained from each chromosome, we then removed the contribution of Xa by subtracting half of the total read-normalized Xa signal obtained in SSCs from the read-normalized FGSC dataset. We used the insulation score to calculate the TAD signal, as previously described^19^. We used the public code to calculate the insulation score in Github (matrix2insulation.pl; <https://github.com/dekkerlab/crane-nature-2015>) with the following parameters: -b 2,000,000, -ids 400,000, -im mean).

**GO Term Enrichment Analysis**

GO term enrichment analysis was performed using the DAVID tool (version 6.8) ^27^, with a focus on enriched biological processes (BP). The GO results were displayed by Cytoscape (version 3.5.1)^28^. For the Benjamin-corrected P value, a threshold of less than 0.05 was used for significance.

**Figure Legends**

**Figure S1**. **Validation of Hi-C Data Quality**

The correlation between Hi-C replicates for each cell type according to the normalized interaction frequency at 400-kb resolution. R indicates Pearson’s correlation coefficient.

**Figure S2**. **Overview of Whole-Genome Interaction Frequency Heat Maps across Five Types of Cells**

**Figure S3**. **Relationship between A/B Compartment and Gene Expressions or Histone Modification (H3K27ac, H3K27me3, H3K4me3)**

(A) Expression of genes with A or B compartment status across five cell types (*: p<0.05; **: p <0.01; ***: p<0.001).
(B) Spearman’s correlation between PC1 values and average ChIP-Seq signal of histone modification (H3K27ac, H3K27me3, H3K4me3).
(C) IGV Browser Showed the A/B Compartment Status and the ChIP-Seq Reads of H3K27ac

**Figure S4**. The Switch of A/B Compartment Correlated with Gene Expression and Histone Modification in FGSCs.

(A) Percentage of switching A/B compartment status in FGSCs, compared to other types of cells.
(B) The change of ChIP-Seq signal between FGSCs and other cells (Left: H3K4me3, Right: H3K27ac; *: p<0.05, **: p<0.01, ***: p<0.001, by Wilcoxon’s test).
(C) Display of the GO enrichment of genes. Different background colors represent different biological processes. The color of each gene represents the compartment status in STOs (blue is the A compartment and orange is the B compartment; adjusted p value < 0.01).

**Figure S5**. **TAD Was Stable but Gene Expression Was Activated in the TAD Boundary**

(A) Boxplot of absolute DI across five types of cells. P value was calculated by Kruskal-Wallis test.
(B) Genome-wide average distribution of RNA-seq, H3K27ac, H3K4me3, and H3K27me3 reads around the domain boundaries in other cells.
(C) Gene expression for the genes located at cell-type-specific or common boundaries.
(D) *Dppa4* showed high expression in the domain boundaries of iPSCs.

**Figure S6**. **Association between Chromatin Loops and A/B Compartment or TAD Types**

(A) Normalized Hi-C maps of chromosome 1 showed concordance between the contact frequency of chromatin loops and *Sox17* expression.
(B) Percentage of chromatin loops located in the regions with different A/B compartment status.
(C) Percentage of chromatin loops located in each TAD type.

**Figure S7**. **The Gene Ontology of FGSCs Specific Chromatin Loops**

The enrichment of gene ontology to FGSCs specific chromatin loops. (Left: loops with active promoters (H3K4me3); Right: loops with active enhancers (H3K27ac).

**Figure S8**. **PC1 Values in Autosomes between FGSCs and SSCs**

PC1 values in FGSCs and SSCs (orange represents FGSCs, blue represents SSCs).

**Figure S9**. **Overview of Interaction Frequency Heatmap in GV**

Left: The whole genome Hi-C heatmap in GV; Right: Intra-chromosome heatmap of Chr19.
