## Supplemental Figures for "Three-dimensional Genome Structure Reveals Distinct Chromatin Signatures of Mouse Female Germline Stem Cells During Development"

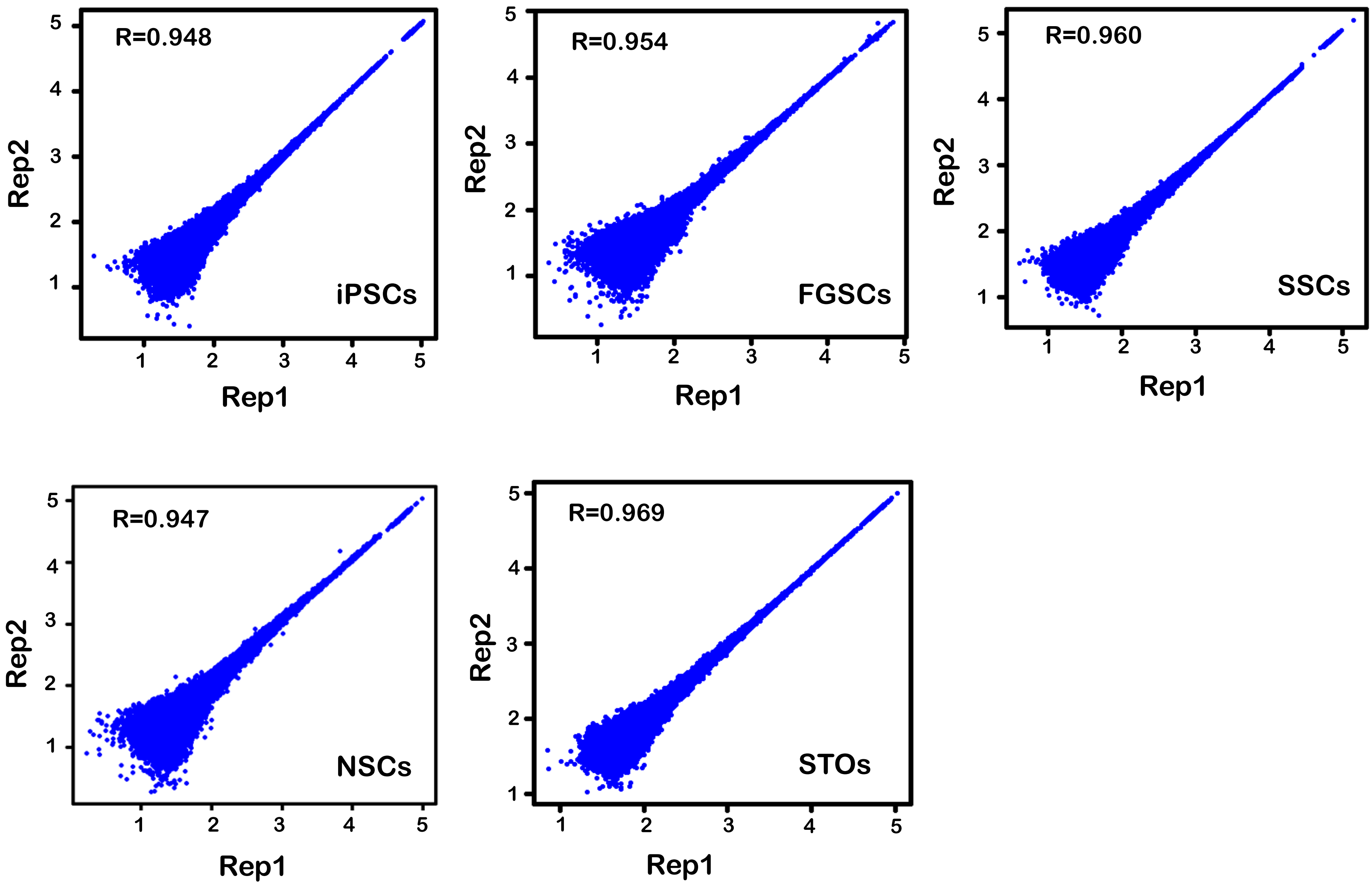
R=0.948
iPSCs
Rep2
Rep1
R=0.954
FGSCs
Rep2
Rep1
R=0.960
SSCs
Rep2
Rep1
R=0.947
NSCs
Rep2
Rep1
R=0.969
STOs
Rep2
Rep1

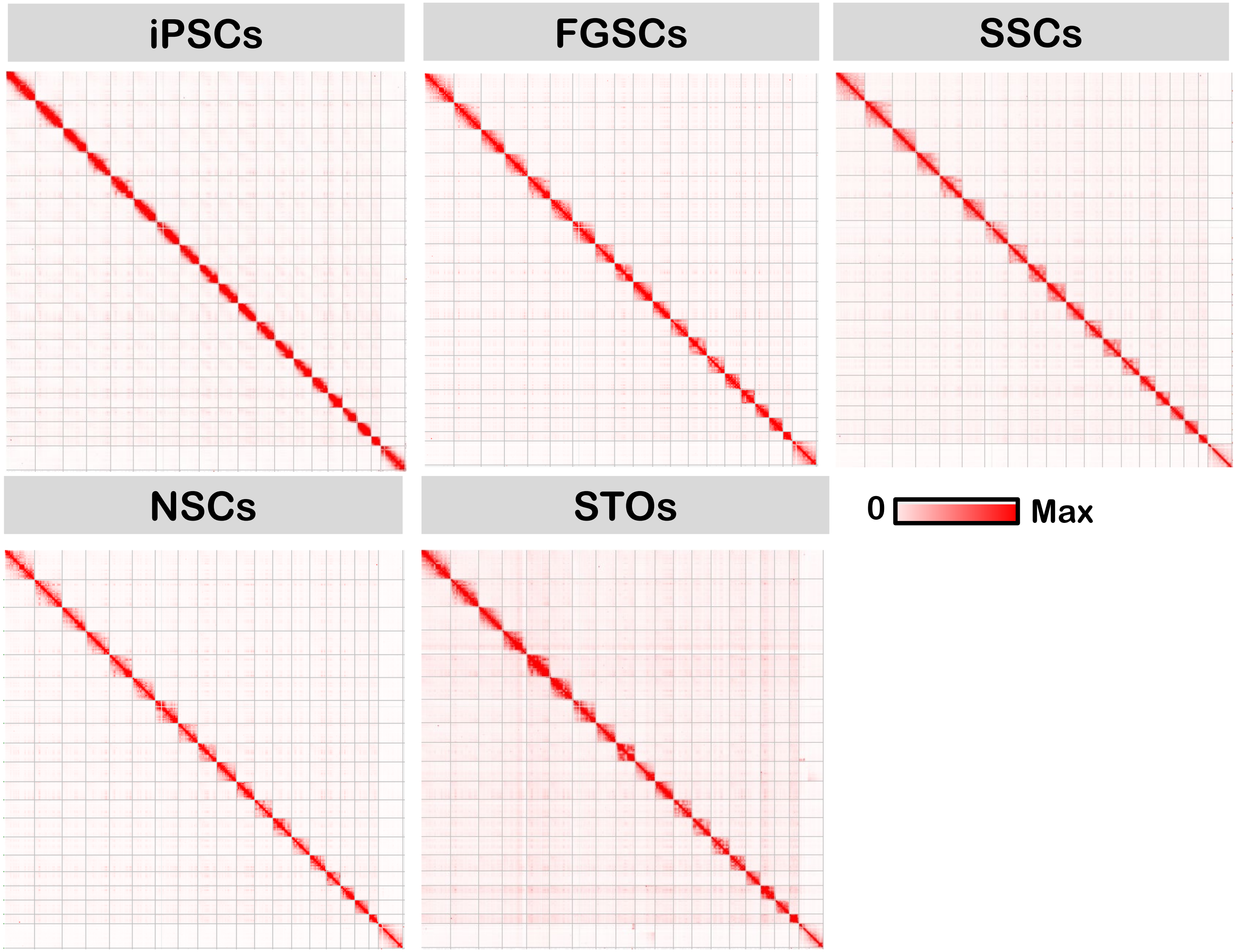

iPSCs
FGSCs
SSCs
NSCs
STOs
0
Max

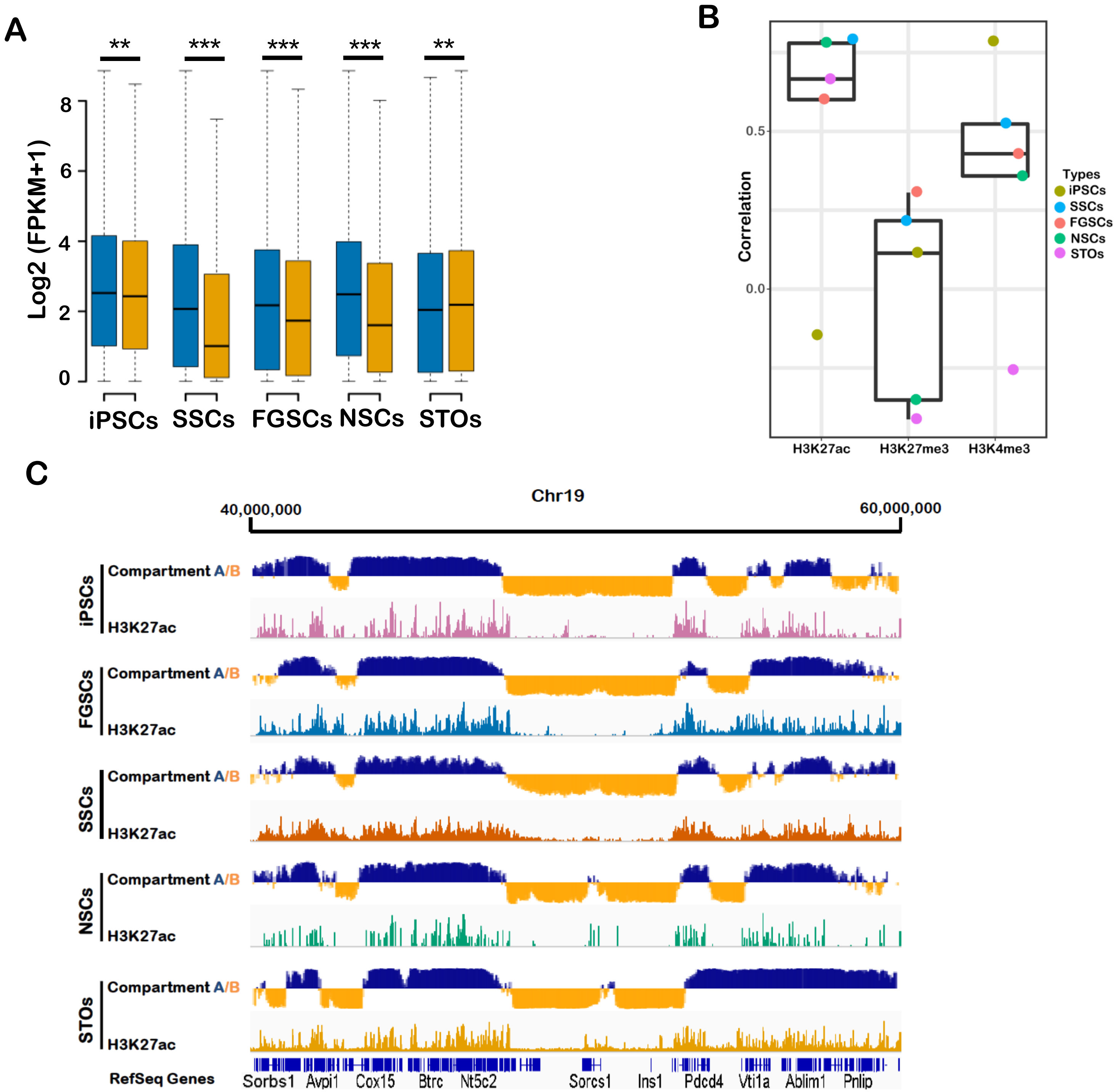

A
**
***
***
***
**
Log2 (FPKM+1)
0
2
4
6
8
iPSCs
SSCs
FGSCs
NSCs
STOs
B
Correlation
0.0
0.5
H3K27ac
H3K27me3
H3K4me3
Types
iPSCs
SSCs
FGSCs
NSCs
STOs
C
Chr19
40,000,000
60,000,000
iPSCs
Compartment A/B
H3K27ac
FGSCs
Compartment A/B
H3K27ac
SSCs
Compartment A/B
H3K27ac
NSCs
Compartment A/B
H3K27ac
STOs
Compartment A/B
H3K27ac
RefSeq Genes
Sorbs1
Avpi1
Cox15
Btrc
Nt5c2
Sorcs1
Ins1
Pdcd4
Vti1a
Ablim1
Pnlip

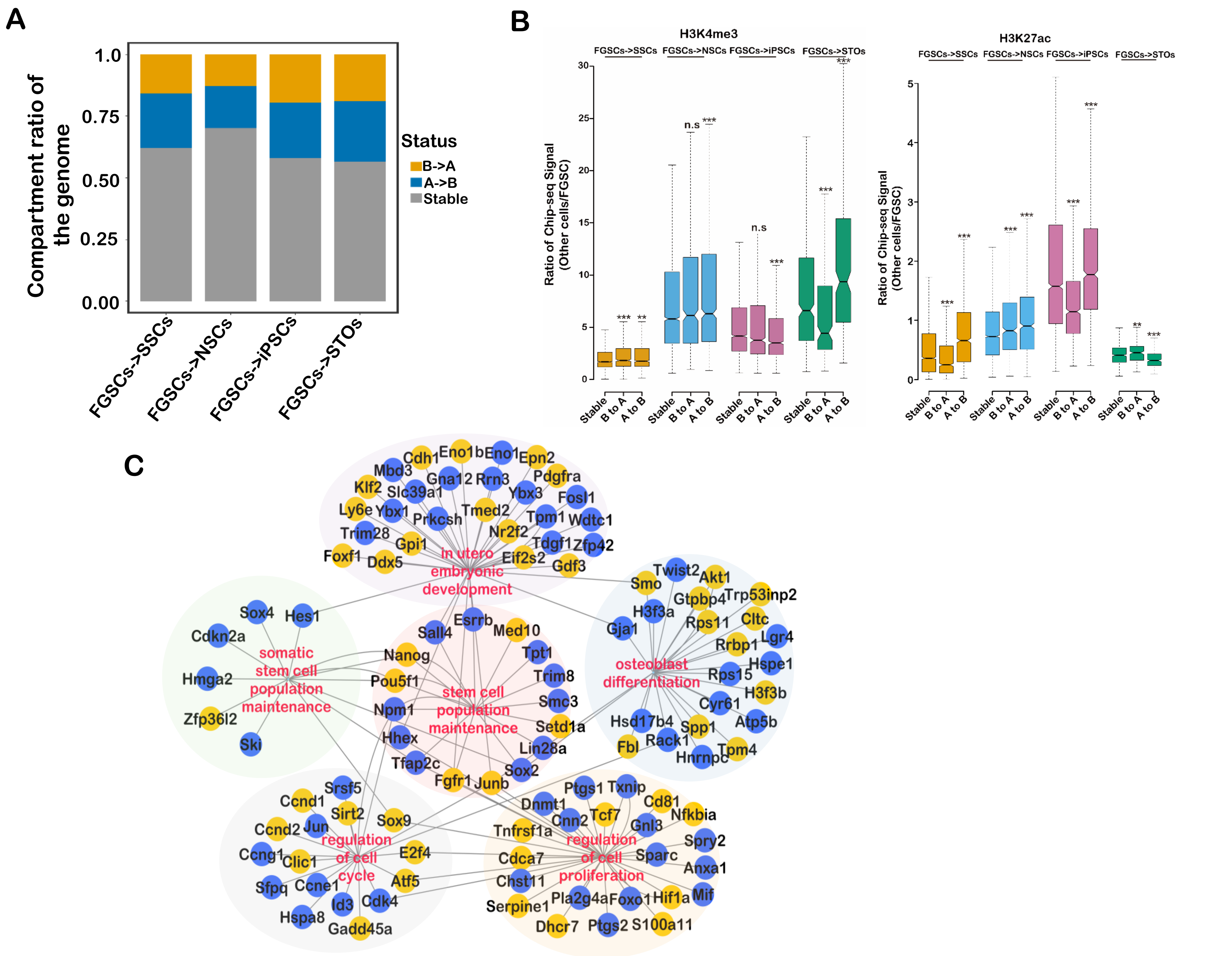
A
Compartment ratio of the genome
1.0
0.75
0.50
0.25
0.00
Status
B->A
A->B
Stable
FGSCs->SSCs
FGSCs->NSCs
FGSCs->iPSCs
FGSCs->STOs
B
H3K4me3
FGSCs->SSCs FGSCs->NSCs FGSCs->iPSCs FGSCs->STOs
Ratio of Chip-seq Signal (Other cells/FGSC)
30
25
20
15
10
5
0
n.s
***
**
Stable
B to A
A to B
H3K27ac
FGSCs->SSCs FGSCs->NSCs FGSCs->iPSCs FGSCs->STOs
Ratio of Chip-seq Signal (Other cells/FGSC)
5
4
3
2
1
0
C
in utero embryonic development
Cdh1 Eno1b Eno1 Epn2 Mbd3 Gna12 Rrn3 Pdgfra Klf2 Slc39a1 Ybx3 Fosl1 Ly6e Ybx1 Prkcsh Tmed2 Tpm1 Wdtc1 Trim28 Nr2f2 Gpi1 Tdgf1 Zfp42 Foxf1 Ddx5 Eif2s2 Gdf3
somatic stem cell population maintenance
Sox4 Hes1 Cdkn2a Hmga2 Zfp36l2 Ski
stem cell population maintenance
Esrrb Sall4 Med10 Nanog Tpt1 Trim8 Pou5f1 Smc3 Npm1 Setd1a Hhex Lin28a Tfap2c Sox2 Fgfr1 Junb
osteoblast differentiation
Twist2 Smo Akt1 Gtpbp4 Trp53inp2 H3f3a Gja1 Rps11 Cltc Rrbp1 Lgr4 Hspe1 Rps15 H3f3b Cyr61 Hsd17b4 Atp5b Spp1 Fbl Rack1 Hnrnpc Tpm4
regulation of cell cycle
Srsf5 Ccnd1 Sirt2 Sox9 Ccnd2 Jun E2f4 Ccng1 Clic1 Atf5 Sfpq Ccne1 Id3 Cdk4 Hspa8 Gadd45a
regulation of cell proliferation
Ptgs1 Txnip Dnmt1 Cd81 Cnn2 Tcf7 Gnl3 Nfkbia Tnfrsf1a Spry2 Cdca7 Sparc Anxa1 Chst11 Mif Pla2g4a Foxo1 Hif1a Serpine1 Dhcr7 Ptgs2 S100a11

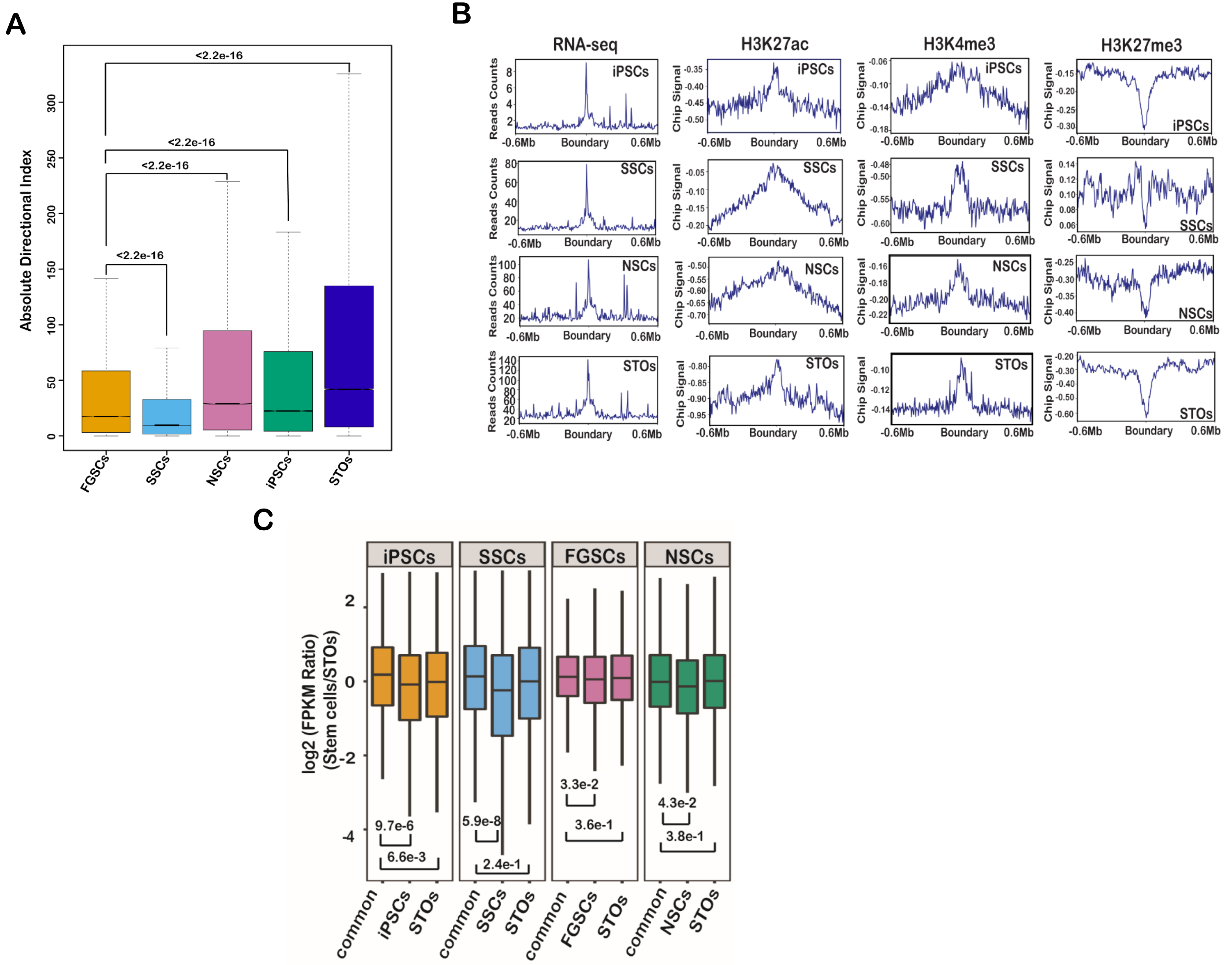
A
Absolute Directional Index
0
50
100
150
200
250
300
<2.2e-16
<2.2e-16
<2.2e-16
<2.2e-16
<2.2e-16
FGSCs
SSCs
NSCs
iPSCs
STOs
B
RNA-seq
H3K27ac
H3K4me3
H3K27me3
Reads Counts
Chip Signal
iPSCs
SSCs
NSCs
STOs
-0.6Mb
Boundary
0.6Mb
C
iPSCs
SSCs
FGSCs
NSCs
log2 (FPKM Ratio)
(Stem cells/STOs)
2
0
-2
-4
9.7e-6
6.6e-3
5.9e-8
2.4e-1
3.3e-2
3.6e-1
4.3e-2
3.8e-1
common
iPSCs
STOs
common
SSCs
STOs
common
FGSCs
STOs
common
NSCs
STOs

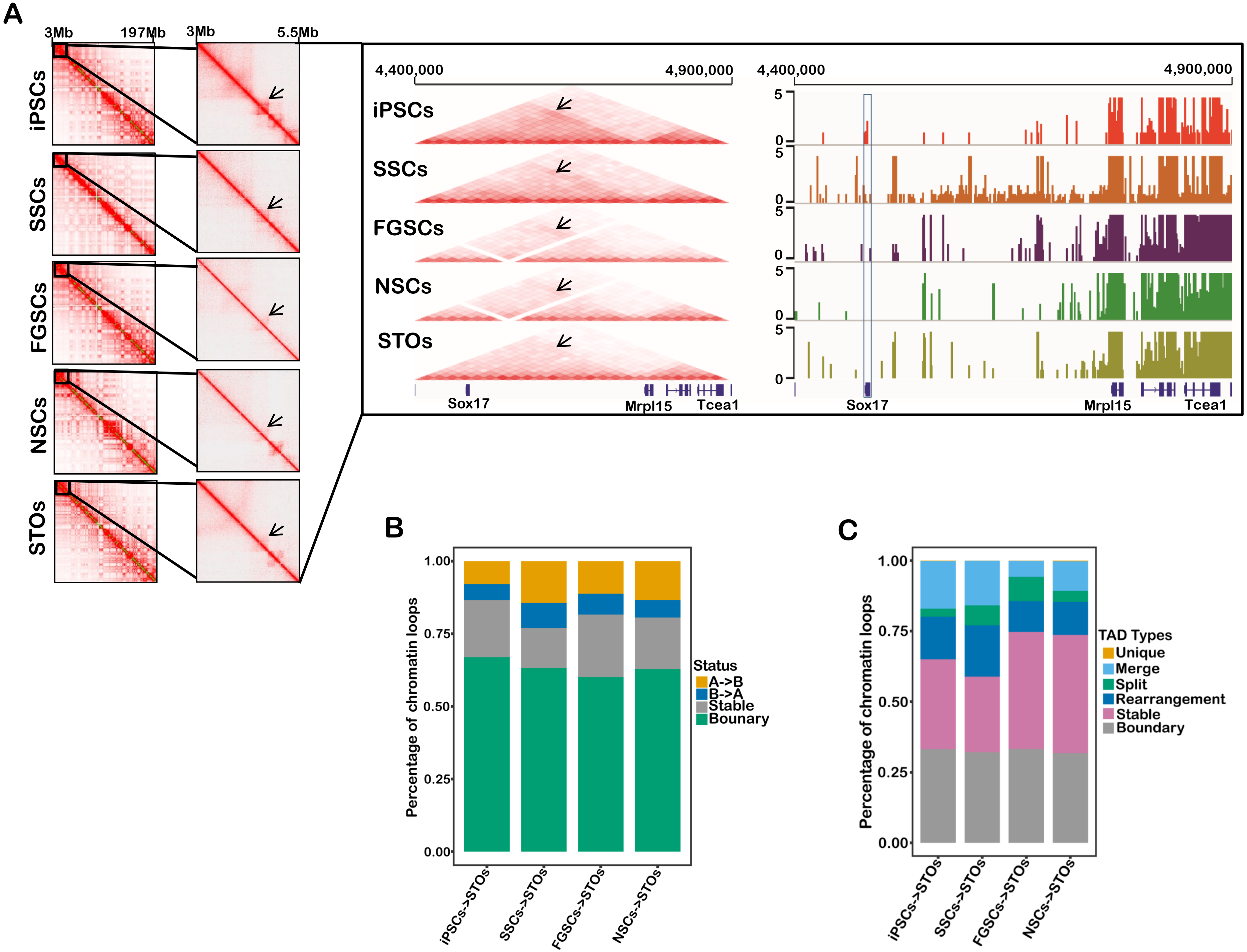

A
3Mb
197Mb
3Mb
5.5Mb
iPSCs
SSCs
FGSCs
NSCs
STOs
4,400,000
4,900,000
4,400,000
4,900,000
5
0
Sox17
Mrpl15
Tcea1
B
Percentage of chromatin loops
1.00
0.75
0.50
0.25
0.00
Status
A->B
B->A
Stable
Bounary
iPSCs->STOs
SSCs->STOs
FGSCs->STOs
NSCs->STOs
C
Percentage of chromatin loops
TAD Types
Unique
Merge
Split
Rearrangement
Stable
Boundary

### The Enrichment of Gene Ontology

#### Active Promoters (H3K4me3)

**Cluster 1**

sensory perception of chemical stimulus
G protein-coupled receptor signaling pathway
sensory perception of smell
nervous system process

**Cluster 2**

regulation of macromolecule metabolic
regulation of metabolic process
regulation of gene expression
developmental process

**Cluster 3**

cellular metabolic process
macromolecule modification
negative regulation of biological process
negative regulation of cellular process
macromolecule modification
organic substance biosynthetic process
biosynthetic process
protein localization
cellular localization

**Cluster 4**

male sex differentiation
sex differentiation
development of primary sexual characteristics
sexual reproduction
regulation of reproductive process
reproductive structure development
reproductive system development

#### Active Enhancers (H3K27ac)

**Cluster 1**

sensory perception of smell
cellular metabolic process
cellular component organization or biogenesis
negative regulation of cellular process
sensory perception of chemical stimulus
G protein-coupled receptor signaling pathway

**Cluster 2**

sex differentiation
development of primary sexual characteristics
sex differentiation
male gonad development
reproductive structure development

**Cluster 3**

nervous system process
system process
nervous system development
cellular component organization or biogenesis
cellular metabolic process
intracellular transport
multicellular organism development
protein transport
regulation of protein modification process

**Cluster 4**

reproductive structure development
reproductive system development
reproductive process
embryo development
tube development
embryonic organ development

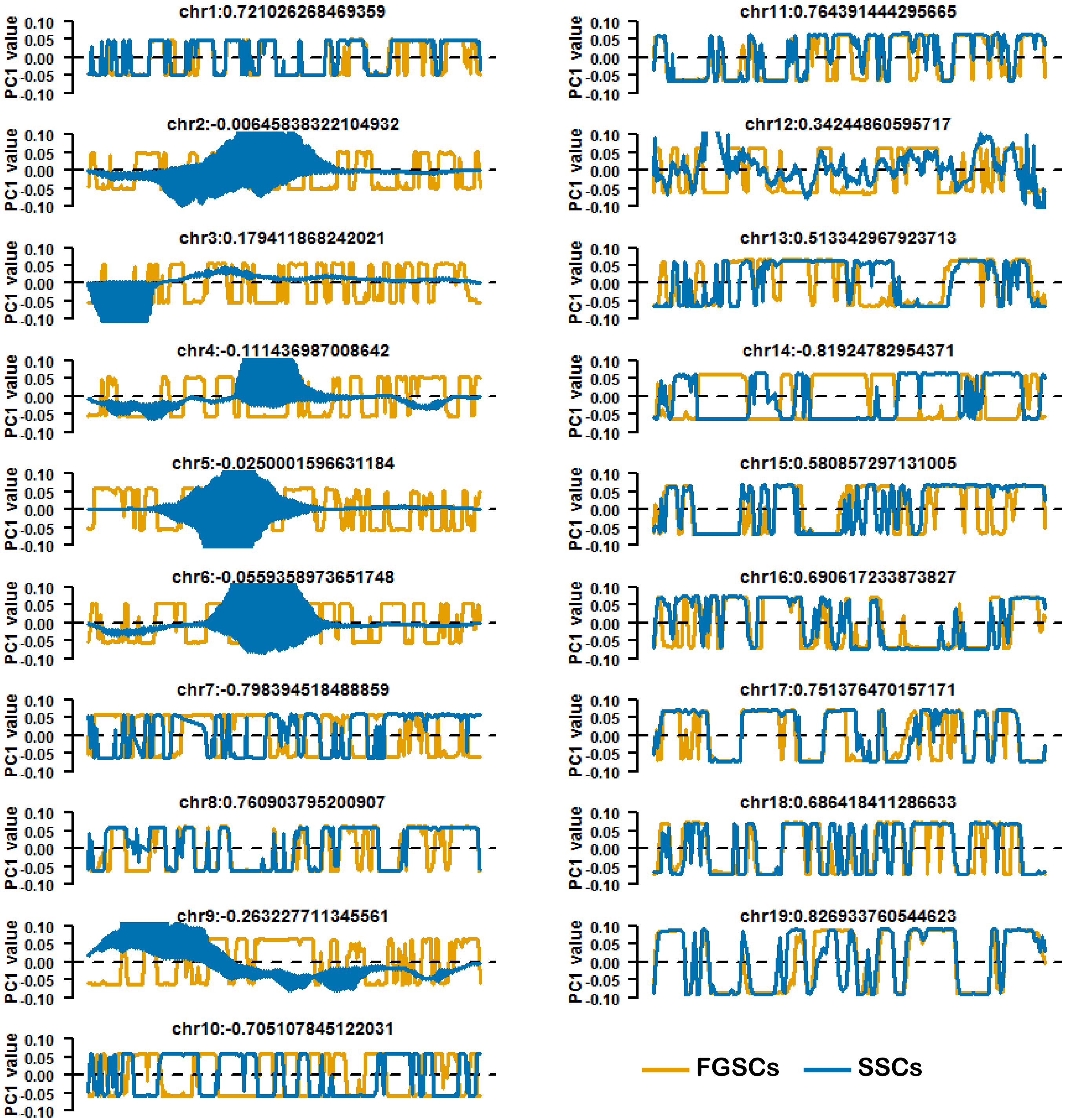
chr1:0.721026268469359
chr2:-0.00645838322104932
chr3:0.179411868242021
chr4:-0.111436987008642
chr5:-0.0250001596631184
chr6:-0.0559358973651748
chr7:-0.798394518488859
chr8:0.760903795200907
chr9:-0.263227711345561
chr10:-0.705107845122031
chr11:0.764391444295665
chr12:0.34244860595717
chr13:0.513342967923713
chr14:-0.81924782954371
chr15:0.580857297131005
chr16:0.690617233873827
chr17:0.751376470157171
chr18:0.686418411286633
chr19:0.826933760544623
PC1 value
0.10
0.05
0.00
-0.05
-0.10
FGSCs
SSCs

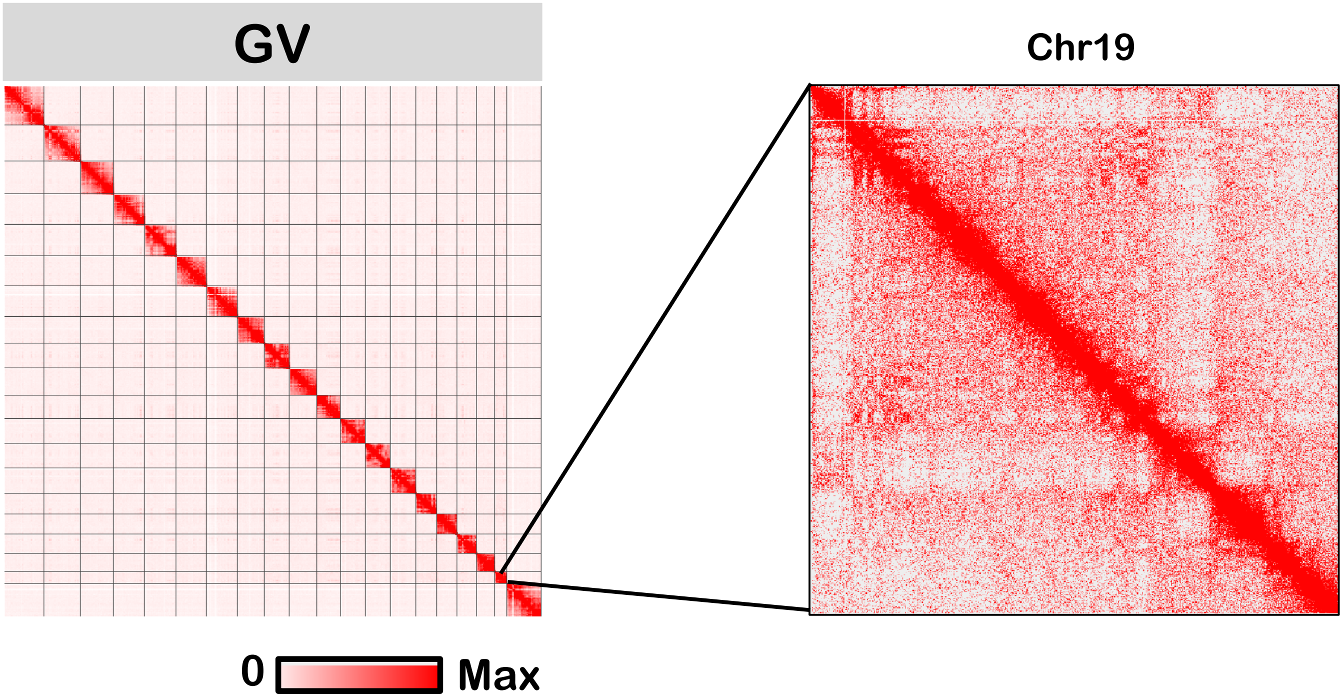
GV
Chr19
0
Max
